# Single nuclei transcriptomics of muscle reveals intra-muscular cell dynamics linked to dystrophin loss and rescue

**DOI:** 10.1101/2022.05.31.494197

**Authors:** Deirdre D. Scripture-Adams, Kevin N. Chesmore, Florian Barthélémy, Richard T. Wang, Shirley Nieves-Rodriguez, Derek W. Wang, Ekaterina I. Mokhonova, Emilie D. Douine, Jijun Wan, Isaiah Little, Laura N. Rabichow, Stanley F. Nelson, M. Carrie Miceli

## Abstract

In Duchenne muscular dystrophy, dystrophin loss leads to chronic muscle damage, dysregulation of repair, fibro-fatty replacement, and weakness. We develop methodology to efficiently isolate individual nuclei from frozen skeletal muscle, allowing single nuclei sequencing of irreplaceable archival samples from small samples. We apply this method to identify cell and gene expression dynamics within human DMD and *mdx* mouse muscle, characterizing treatment effects of dystrophin rescue by exon skipping therapy at single nuclei resolution. *DMD* exon 23 skipping events are directly observed and increased in myonuclei from treated mice. We describe partial rescue of type IIa and IIx myofibers, expansion of a novel MDSC-like myeloid population, recovery of repair/remodeling M2-macrophage, and repression of inflammatory POSTN1+ fibroblasts in response to exon skipping and partial dystrophin restoration. Use of this method enables exploration of cellular and transcriptomic mechanisms of dystrophin loss and repair.

## Introduction

Duchenne muscular dystrophy (DMD) is caused by loss of function mutations in *DMD*, encoding dystrophin. Lack of dystrophin leads to contraction-induced myofiber injury, immune infiltration (*1*) and, ultimately, replacement of myofibers by fat and fibrosis. Loss of myofiber promotes progressive skeletal, diaphragmatic and cardiac muscle weakness and premature death. In healthy individuals, acute muscle injury triggers tightly coordinated immune cell infiltration and fibroblast expansion, essential for clearing damaged tissue and guiding satellite cell activation, differentiation, and muscle regeneration. Tissue remodeling occurs locally, at the site of injury, and resolves once the muscle is repaired. In DMD, chronic and heterogeneous myofiber damage results in dysregulation of immune- and fibroblast-coordinated muscle repair. Dystrophin replacement/repair strategies, including exon skipping, nonsense mutation readthrough and micro-dystrophin gene therapy (*2*), demonstrate some evidence of dystrophin replacement, muscle repair and clinical benefit (*3*). It remains unclear which cellular and molecular aspects of dysregulated muscle tissue remodeling are reversible by dystrophin rescue in the context of ongoing disease.

To demonstrate our sample sparing single nuclei sequencing technique, we used eleven archived frozen tibialis anterior samples from our published cohort including four 6 month old *mdx* mice, four *mdx* mice treated for 6 months with weekly exon23 directed phosphoramidite morpholino oligomer (e23AON), and three genetically matched C57BL/10ScSnJ controls. We have previously validated partial dystrophin rescue by western blot (ranging from 2-12% of wildtype levels) and reversal of pathology in *mdx* mice treated with e23AON in this cohort(*4*). Similar findings using RT-PCR have also been reported in a ΔEx51 transgenic mouse model(*5*). Recently, gene expression has been observed within individual cells or nuclei within heterogenous tissues including human (*6, 7*), and mouse muscle(*8–20*) but most methods require whole cells from fresh tissue. Whole cell methods are unable to examine myofibers due to size constraints and could not explore the diversity of nuclei the within multinucleated cells. Here we describe a novel method to purify nuclei from small numbers of cryotome sections of frozen muscle (~3mg) for high quality single nuclei RNA sequencing (snRNAseq). We present here the largest single cell/nuclei dataset of dystrophic muscle published to date. Using this new single nuclei dataset, we identify cell regulatory processes involved in muscle degeneration, pathology, and repair in a DMD mouse model. We demonstrate that morpholino induced dystrophin rescue partially reverses pathologic cell population and gene expression changes of myolineage cells, fibroblasts and immune cell populations. Further, we apply the method to the study of archived frozen human muscle biopsies from healthy and DMD patients, producing the first snRNAseq report of human dystrophic muscle, and revealing substantial cellular heterogeneity and aberrant gene expression in human DMD.

## Results

To simultaneously assess gene expression of mono- and multi-nucleated cells within skeletal muscle, we developed a method of nuclei extraction from small quantities of frozen muscle tissue sections (Fig. 1a). Frozen tissues were disaggregated, and intact nuclei were sorted away from debris using flow cytometry. Even with the diversity of cell types within skeletal muscle, a single population of intact nuclei was identified with a size range from 7-10μm (Fig. 1b). Quality and quantity of purified nuclei were verified (Fig. 1b) with yields from 3mg of muscle ranging from 6,000-100,000 nuclei.

**Figure 1.**
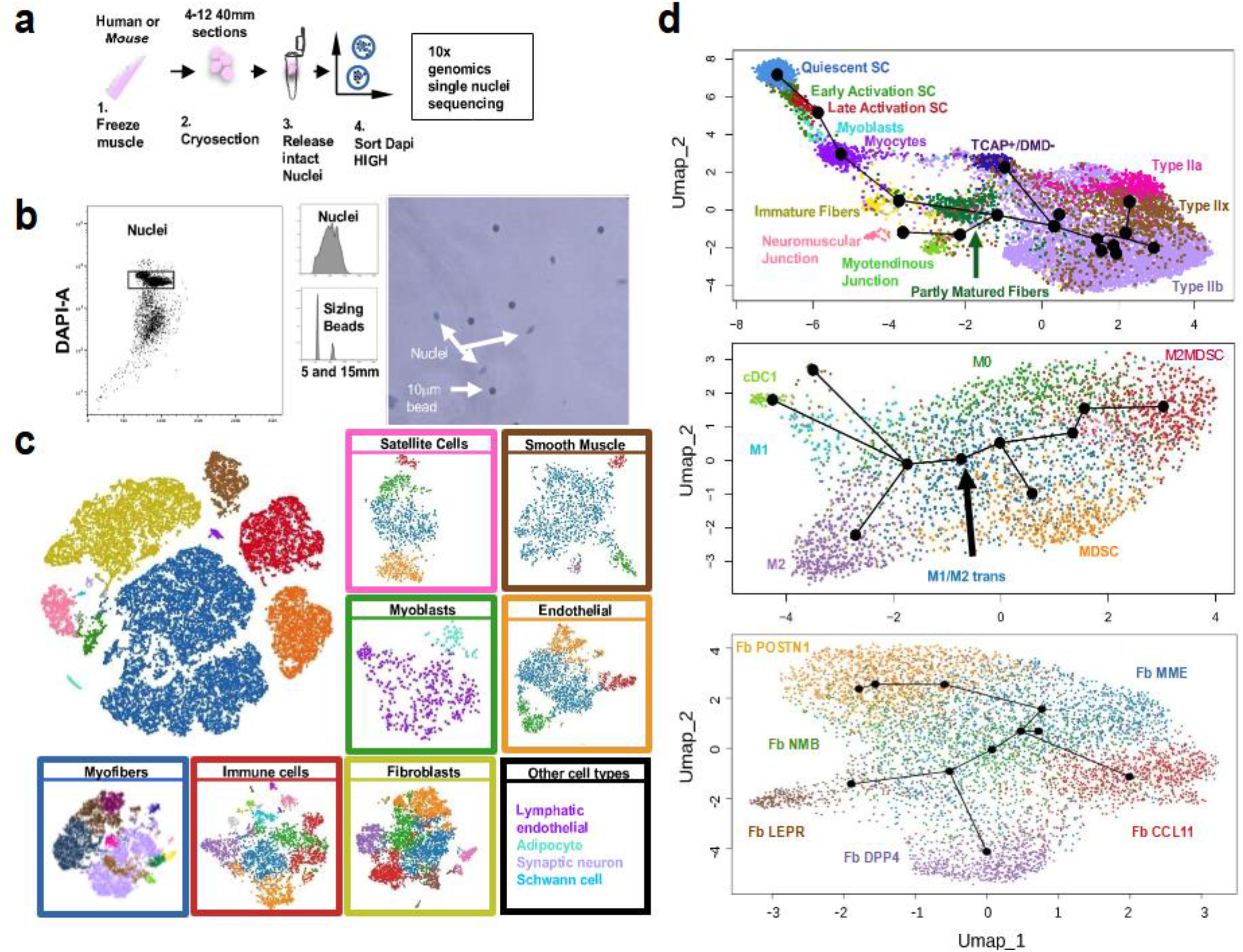
Isolation of single nuclei from frozen muscle and identification and characterization of cell types found in murine muscle (**a**) Visual depiction of nuclei purification for snRNA sequencing. (**b**) Representative flow cytometric sorting of nuclei gated on DAPI staining (left panel), demonstration of forward scatter comparison to beads (middle panels), and size and morphology confirmation by visual microscopy (right panel). (**c**) Major cell types were identified (top left panel) and each major cluster was re-clustered separately to identify subsets within each group (box colors surrounding subcluster correspond to colors in top left panel). Cell types with fewer members are listed in “Other cell types” panel, text color reflects the color of the population in the top left panel. (**d**) Lineage tracing analysis of subpopulations identifies relatedness: (**Top**) Satellite cells/Myoblasts/Myofiber, (**Middle**): Monocyte/Macrophage/DC, (**Bottom**): Fibroblast populations. Text in plots mark each subcluster population adjacent to each population and are colored to match the sub-cluster identity of panel C.

Up to 20,000 nuclei/sample were loaded on to the 10X Genomics Chromium™ Controller Single Cell Sequencing System; ~35% of loaded nuclei resulted in viable libraries. The resulting libraries showed a single peak at ~450bp and were sequenced to about 200 million paired-end reads per library, generating ~40,000 reads per nucleus. Eleven mouse tibialis anterior (TA) muscle samples yielded 34,833 single nuclei libraries, averaging 3,167 nuclei libraries per sample. After normalizing sample read depth, samples averaged 39,334 reads per nucleus with 90% sequence saturation, resulting in 836 genes and 1,371 UMI per nucleus. Nuclei with <200 genes per nucleus and the 4,960 predicted doublets identified by DoubletFinder (*21*) were removed from the dataset. The remaining mouse dataset contained 29,873 nuclei (9,967 nuclei from *mdx*, 8,411 from *mdx* e23AON, and 11,495 from WT muscle) with 846 genes per cell and 1396 UMI per nucleus. Five human vastus lateralis (VL) muscle biopsies generated a total of 7,549 nuclei (4,906 from DMD and 2,643 from healthy individuals) with a median of 655 genes and 993 UMI per nucleus.

### Transcriptional profiling identifies 45 discrete populations within healthy and dystrophic mouse skeletal muscle

We performed TSNE analysis combining all 29,873 mouse single nuclei data to enable robust cell type identification. Nuclei from 11 major cell types were identified based on similarity of gene expression including: satellite cells, myoblasts, myofibers, immune cells, fibroblast/fibro-adipogenic progenitors (FAP), smooth muscle, endothelial cells, adipocytes, neurons, Schwann cells (Fig. 1c). These main populations were further refined by sub-clustering of nuclei, ultimately identifying 45 discrete clusters highlighting cellular heterogeneity of muscle tissue (Fig. 1c).

We identified 13 nuclei clusters among skeletal muscle lineage populations (Fig. 1c, Fig. 2a, Table 1) reflecting nuclei characteristic of mononuclear cells including satellite cells (quiescent, early, and late activation stage), myoblasts, myocytes, as well as multi-nucleated cells characteristic of immature and partially mature fibers, type IIa, IIx, and IIb myofibers, as well as nuclei indicative of specialized functions within myofibers including neuromuscular junction (NMJ) nuclei, myotendinous junction (MTJ) nuclei, as well as Tcap nuclei, a previously undescribed myonuclei type strongly expressing *Tcap* (encoding titin cap), but not *Dmd*. No type I myofiber nuclei were observed, as expected for mouse TA. The smooth muscle compartment included nuclei from pericytes (*22*), vascular smooth muscle cell type1 and type2 (*22*), (*23*) (Fig. 1c, Fig. 2e, Table 1), and an uncategorized smooth muscle cell type (SM). Within the endothelial compartment, we identified 4 subpopulations, including tip-like, stalk-like (*24*), angiogenic, and muscle-derived endothelial cells (Fig. 1c, Fig. 2e, Table 1).

**Figure 2.**
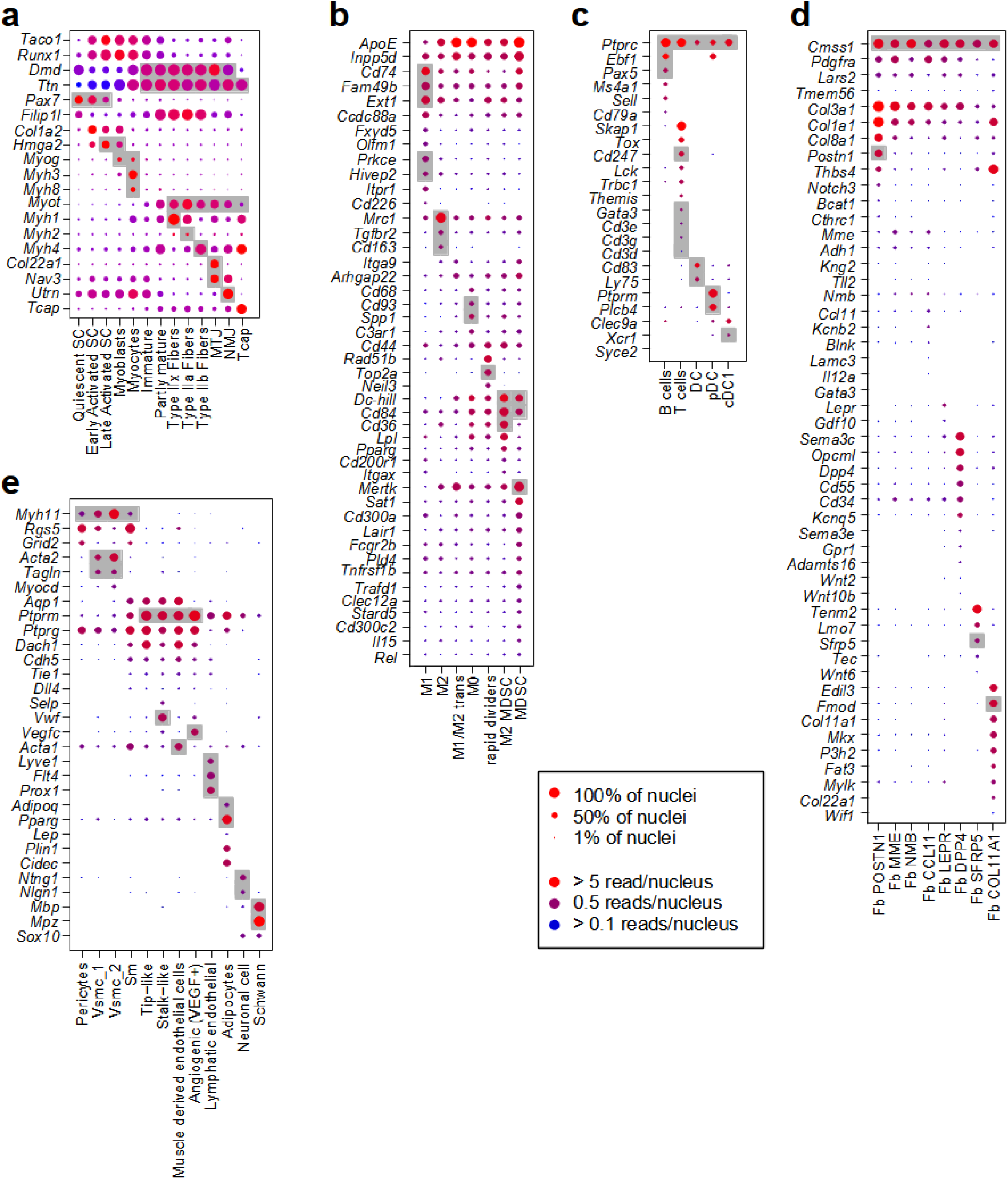
Cell type identity based on expression of significantly expressed genes across major cell types Gene expression differences between subtypes of major cell classes, (**a**) Myolineage, (**b**) Macrophage, (**c**) other immune cells, (**d**) Fibroblasts, (**e**) other cell types. Subtype names indicated on x-axis, gene names indicated on y-axis. Log2(Average gene expression) for each subtype show by Blue-Red gradient. Percentage of cells expressing genes from 0% to 100% indicated by dot size. Gray background indicates known markers of the respective cell type on x-axis. Marker genes are listed in Figure S2, Table S1a, b, Data file S1.

**Table 1.**
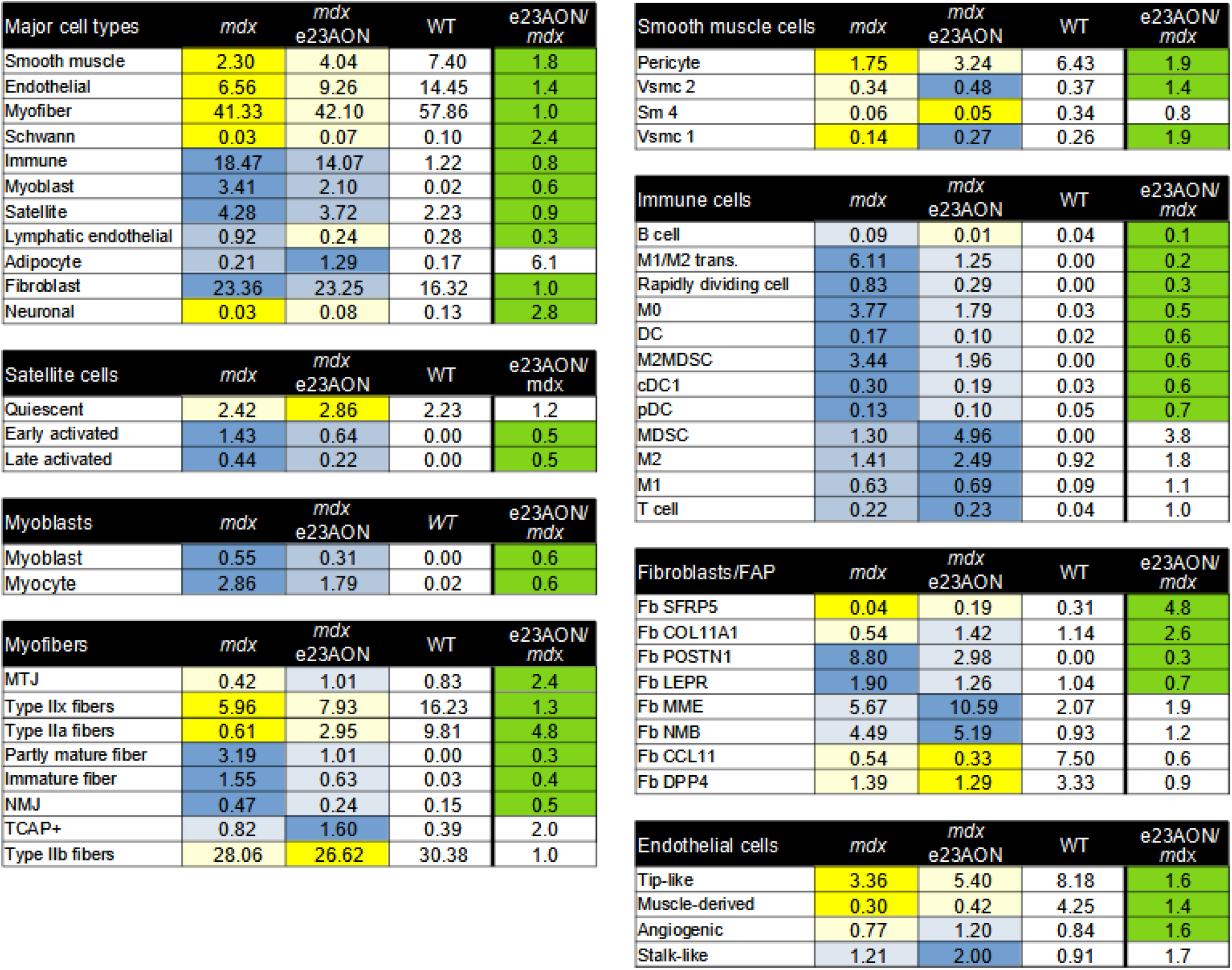
Frequencies of intra-muscular cell populations in WT and *mdx* mice and partial normalization of most populations in response to e23AON treatment of *mdx*. Percentages of each cell population for each pooled treatment condition, *mdx, mdx* e23AON, WT. Following the major cell types, cell types are further broken down to their respective subclusters. Blue = increase in percentage relative to WT, Yellow = decreased in percentages relative to WT. Rightmost column shows the effect of e23AON treatment relative to *mdx*, green denotes that cell numbers of *mdx* treated with e23AON recovered a more WT percentage.

Eight fibroblasts/FAP subtypes were identified and named based on differentially expressed genes (Datafile S1) with potentially relevant function: Fb CCL11, Fb DPP4, Fb POSTN1, Fb SFRP5, Fb COL11a1, Fb LEPR, Fb MME, Fb NMB (Fig. 1c, Fig. 2d, Table 1). Several populations share features with fibroblast/FAP populations reported previously at the single cell level (*25*). Twelve different CD45+ immune cell types were identified; B cells, T cells, DC, cDC, pDC, M1-macrophage, M2-macrophage, M1/M2 transitional macrophage, two less well differentiated macrophage cell types, termed M0 and rapidly dividing, and two previously unrecognized cell types expressing several markers of myeloid derived suppressor cells and negative regulators of inflammation, here referred to as MDSC, and M2MDSC (Fig. 1c, Fig. 2b, c, Table 1). Definitive identification of these myeloid populations as MDSC will require functional assessment of these populations(*26*) (Fig. 2b and Table S1e, Complete lists of genes differentially expressed between MDSC and M2MDSC populations can be found in Data file S2).

### Lineage trajectory analysis elucidates developmental and transcriptional relationships between cell types and confirms cell identity

To quantify relatedness of identified cell types, we performed pseudotime lineage tracing analysis on myogenic and mature muscle fiber, immune, and fibroblast lineage nuclei (*27*). This analysis showed a detailed developmental trajectory of myolineage cells, consistent with our annotated cell-type identifications. This linear trajectory created solely by gene expression similarity was consistent with established myofiber development from quiescent SC though to myoblast, myocyte, to mature myofiber subtypes, including type IIa, IIx, and IIb myonuclei, NMJ, MTJ, and Tcap (Fig. 1d, top) (*28*).

Myeloid immune cells and fibroblasts are known for their plasticity and responsiveness to environmental cues via and activation of gene programs and differentiation (*29*). Lineage trajectory analysis of intramuscular myeloid nuclei indicate a high degree of relatedness, despite expression profiles reflecting distinct functionality, raising the possibility of trans-differentiation of one subset into an another in the context of differing muscle micro-environments (Fig. 1d, middle). Consistent with reports that MDSCs arise from aberrant early myelopoiesis, the MDSC-like population was more distantly related to differentiated M1 and M2 effector populations, while the M1/M2 transitional cells were equally related to MDSC,M1, and M2, suggesting that M1/M2 are developmental intermediates (Fig. 1d, middle).

Lineage tracing of the fibroblast subpopulations revealed a spectrum of diverse but highly related fibroblasts. Fb CCL11 and Fb DPP4 were least related to Fb POSTN1, with Fb NMB and Fb MME representing potential developmental intermediates (Fig. 1d, lower). Fb SFRP5 and Fb COL11A1, were not represented in Fig. 1d (lower), as they cluster as distinct entities and are less related to the other fibroblast populations (Fig. 1c, Fibroblast Pink and Grey subclusters). Fb COL11A1 is likely perimysial cells (*25*) present at the myotendinous junction (*Col22a1, Mkx*). Fb SFRP5 also expressed *Tenm2*, found at sites of muscle attachment (*30*).

### Exon 23 skipped *Dmd* mRNA is observed at single nuclei resolution

In order to gain insight into mechanisms of degeneration and repair in dystrophic muscle, we compared cell and gene expression profiles of nuclei isolated from frozen muscles from WT control mice (n=3), *mdx* dystrophic mice (n=4), and *mdx* mice treated with e23AON (n=4) (*4*). Systemic delivery of e23AON in *mdx* mice causes a portion of mature *Dmd* mRNAs to delete exon 23 (e23), removing the stop codon in e23 creates an in-frame mRNA that is translated into an internally deleted protein (*4*). We retrieved all reads from snRNAseq fastq files that map to *Dmd* e22-e24 junctions (indicative of e23 skipping) and map their expression along the myolineage developmental trajectory (Fig. 1d, top). While e22-e23 junctions were detected as early as the myocyte phase, e22-e24 junctions were only observed in mature myonuclei (Fig. 3a). As expected, both junctions were rare in our dataset (1.22% of myonuclei contain reads spanning e22-e23, and 0.08% of myonuclei contain reads spanning e22-e24) and are thus an under-observation of actual exon skipping (*31*). The frequency of e22-e23 junctions was similar in both *mdx* and e23AON-treated *mdx*, however the frequency of e22-e24 junctions was about 10x higher in e23AON-treated *mdx*. This is roughly consistent with dystrophin rescue of between 2% and 12% of WT dystrophin levels previously reported (Fig. 3b, Fig. S3) (*4*).

**Figure 3.**
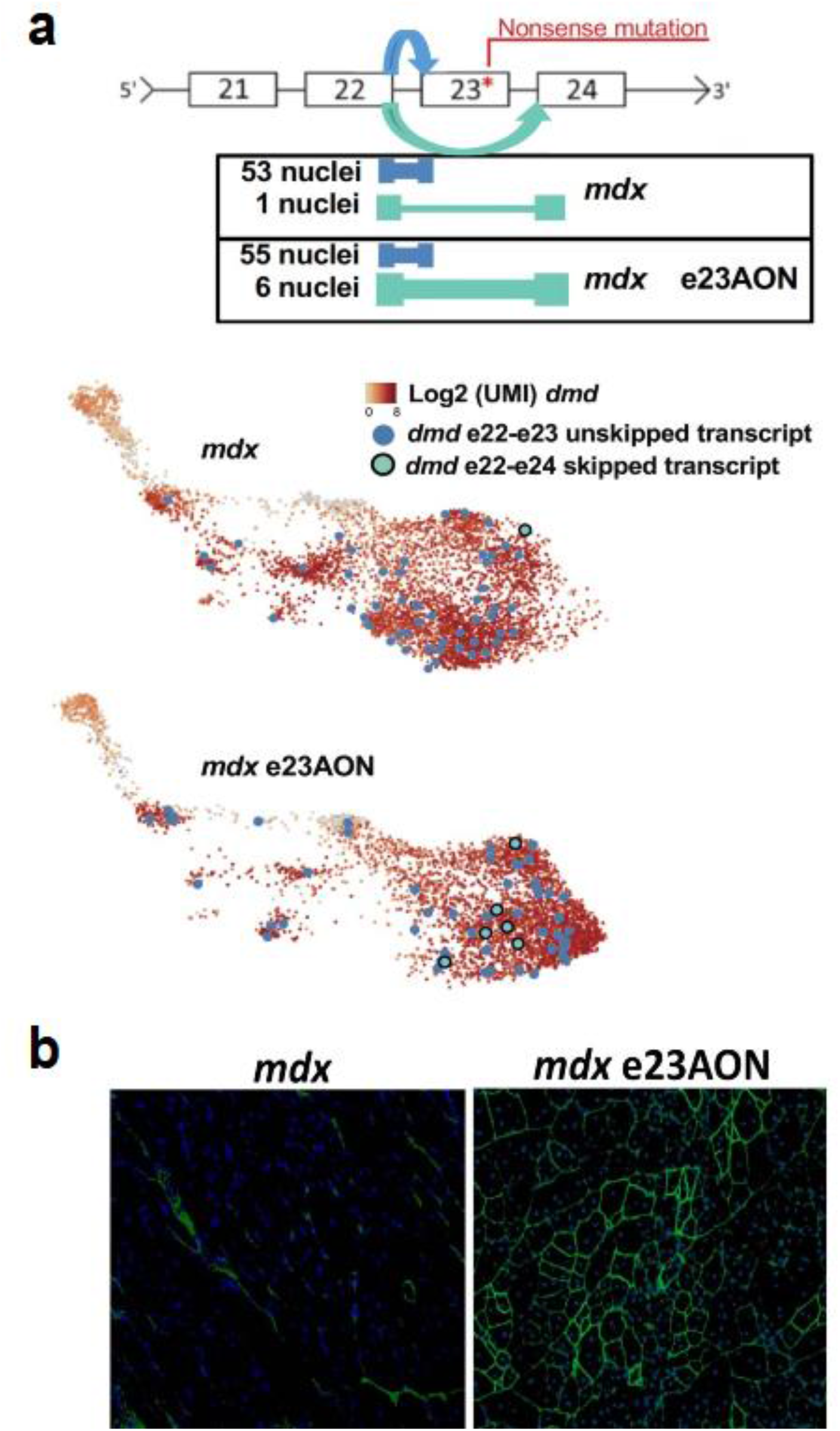
Exon skipping events are mapped to specific nuclei in e23AON-treated *mdx* (**a**) Diagram of *Dmd* expression and e22-e24 *Dmd* exon skipping. Number of nuclei containing *Dmd* e22-e24 (skipped light blue) and e22-e23 (unskipped dark blue) junction reads are shown to the left of the splicing events for each treatment condition. Nuclei where e22-e24 was observed are color coded on the UMAP of myolineage nuclei (from Figure1D) with overall *Dmd abundance shown* (Yellow-Red color scale). (**b**) Dystrophin protein detection (green-FITC) by immunohistochemistry in muscle from *mdx* (left panel) and e23AON-treated *mdx* (right panel). Whole muscle stains shown in Figure S3.

### Muscle remodeling occurs with dystrophin deficiency and partial rescue and is reflected in altered cellular composition

The relative percentages of nuclei from all major intramuscular cell types across WT, *mdx* and *mdx* e23AON mice are shown in Table 1. In WT TA muscle, the majority of nuclei represented mature myofibers (10% type IIa, 16% IIx, and 30% IIb), with few resident quiescent SC (2.2%), anti-inflammatory M2 macrophage (0.9%), fibroblast (16%) and other cell (23%) populations present. *mdx* had a substantial loss of mature myofiber nuclei, predominantly due to selective loss of type IIa (1%) and IIx (6%) myofibers. This is coincident with a hundred-fold expansion of myoblasts, a doubling of satellite cells, a 15-fold increase in immune cells and more modest increases in fibroblasts.

Three subpopulations of satellite cells were identified; quiescent, early activated, and late activated (*28*) (Fig. 4, 1^st^ row). Whereas WT muscle contained only quiescent satellite cells, *mdx* showed expansion of all three satellite cell populations as well as myoblasts and myocytes, reflecting increased regeneration in *mdx muscle*. Furthermore, “quiescent” *mdx* satellite cells cluster more closely to activated satellite cells, suggesting disruption of complete quiescence within *mdx* muscle relative to WT. Dystrophin rescue was associated with an increased percentage of type IIa (3%) and IIx (8%) myonuclei within e23AON-treated *mdx*, accompanied by a reduction in both satellite cells and myoblasts (6%) and the overall immune cell infiltrate (14%), reflecting partial normalization toward WT cell numbers (Fig. 4, 1^st^ and 2^nd^ row, Table 1).

**Figure 4.**
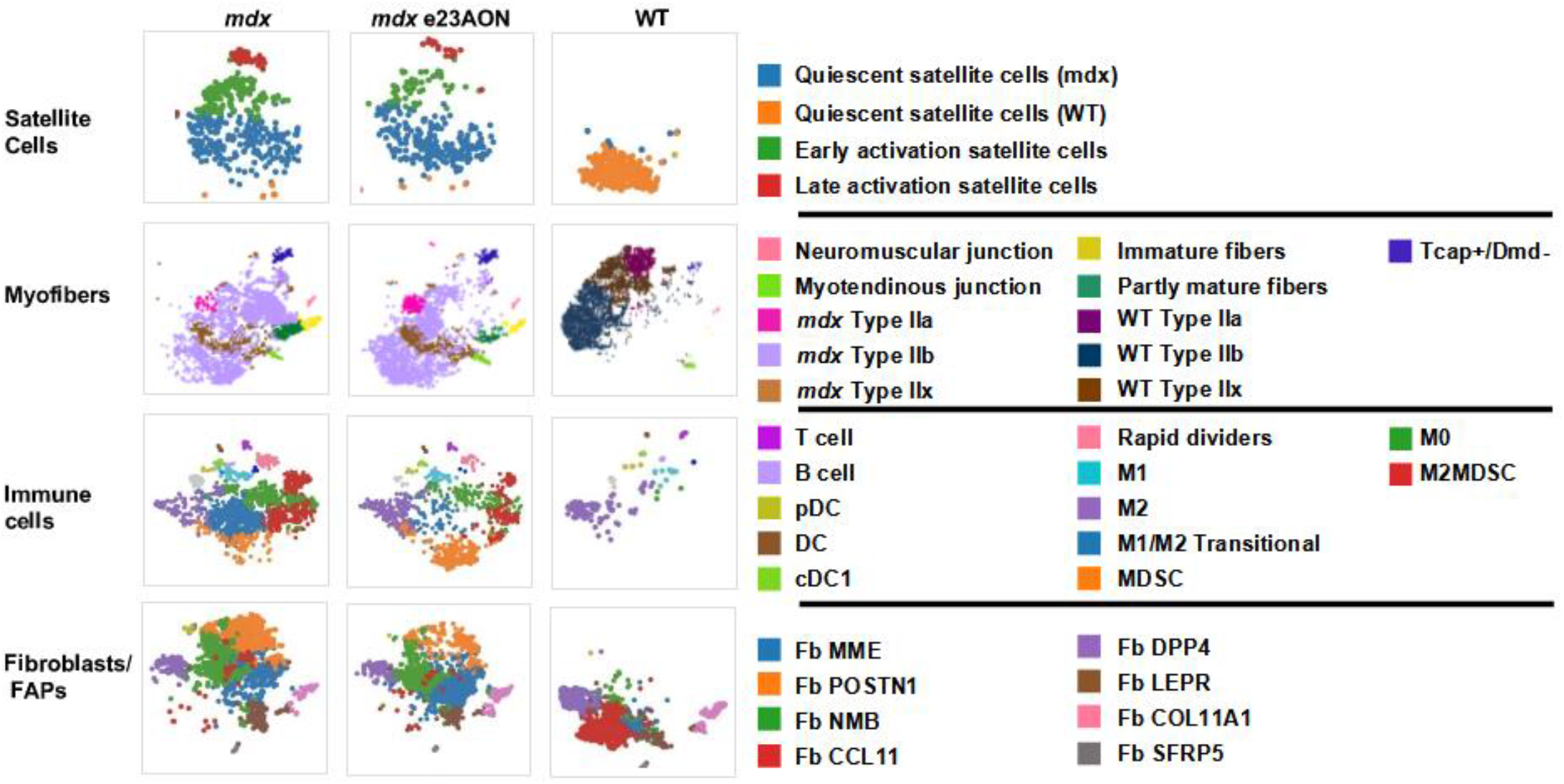
e23AON treatment drive shifts in subcluster identity in *mdx* toward a more WT pattern. TSNE plots for major cell populations across each treatment condition. Sub-clusters identified in Figure1 derived from quiescent satellite cells (**1^st^ row**), myofibers (**2^nd^ row**), CD45+ immune cells (**3^rd^ row**) and fibroblasts/FAP (**4^th^ row**) are separated by treatment condition (*mdx, mdx* e23AON, and WT). Sub-cluster identities are indicated by the color-coded box to the left of the cell subset name, which matches the color of the cluster of cells identified.

Myeloid cells represented the majority of intramuscular immune cells. M2 macrophages were the primary population in WT muscle, representing less than 1% of total nuclei. In *mdx* dystrophic muscle, multiple myeloid lineage cells were expanded and diversified, cumulatively representing 17% of total nuclei (Table 1), including M1-like (M1) (0.63% of total nuclei), M0 (3.78%), M1/M2 transitional cells (6.11%), M2 (1.41), MDSC (1.3%) and M2MDSC (3.44%) in *mdx* muscle (*32*), (*31*). Percentages of M0 (1.79%), M1/M2 transitional cells (1.25%), M2MDSC (1.96%) and B cells (0.01%) were substantially decreased with e23AON treatment, reflecting partial normalization of dystrophic intramuscular immune subsets toward WT composition (Fig. 4, 3^rd^ row, Table 1). e23AON treatment increased the frequencies of some cells with potential immune suppressive/anti-inflammatory properties, MDSC (4.96%) and M2 macrophage (2.49%), driving them even higher than WT frequencies (Fig. 4, 3^rd^ row, Table 1). DC, B and T cell populations were also expanded in *mdx* relative to WT mice (Fig. 4, 3^rd^ row, Table 1).

Within the fibroblast compartment, Fb DPP4 (3.33%) and Fb CCL11(7.5%) were the most common types within WT muscle. Fb POSTN1 was not observed in WT, but it was the dominant type in *mdx* muscle (8.8% of total nuclei). Fb SFRP5, present at low levels in WT, was diminished further in *mdx*. Fb NMB and Fb MME appeared to be intermediate populations and were present at low levels in WT, while expanded in *mdx*. Based on pseudo-lineage tracing and gene expression, we inferred that *mdx* expanded fibroblast populations represent trans-differentiation from resident anti-inflammatory fibroblasts observed in WT muscle, to relatively more proinflammatory/profibrotic fibroblasts (Fig. 4, 4^th^ row, Table 1, Table S1b) (*33*) (*34*) (*35*).

The prominent *mdx* profibrotic Fb POSTN1 decreased with e23AON treatment (8.8% to 2.98%). Fb MME and Fb NMB populations, increased in *mdx* versus WT, expanded to even higher levels in response to e23AON (10.6% for Fb MME and 5.19% for NMB). Based on fibroblast lineage tracing (Fig. 1d, lower), increases in these fibroblast populations may represent the accumulation of intermediates transitioning toward Fb DPP4 and Fb CCl11, the dominant WT fibroblast. Fb LEPR contracts, and Fb SFRP5 and Fb COL11A1 expanded with e23AON treatment, reflecting more WT representation.

### Dystrophin loss and rescue impacts gene expression signatures in multiple cell types

We assessed differential gene expression between *mdx* and WT mice nuclei, and between *mdx* and e23AON-treated *mdx* in each of the 45 identified cell/nuclei clusters. There was a trend that differentially expressed (DE) genes between corresponding populations in *mdx* and *mdx* e23AON, acquired a more ‘WT-like’ expression level with e23AON treatment. This was true for DE genes in myofibers and satellite cells (Fig. 5a), most immune (Fig. S4) and fibroblast (Fig. 6e, Fig. S4), and some other sub-populations (Fig. S4).

**Figure 5.**
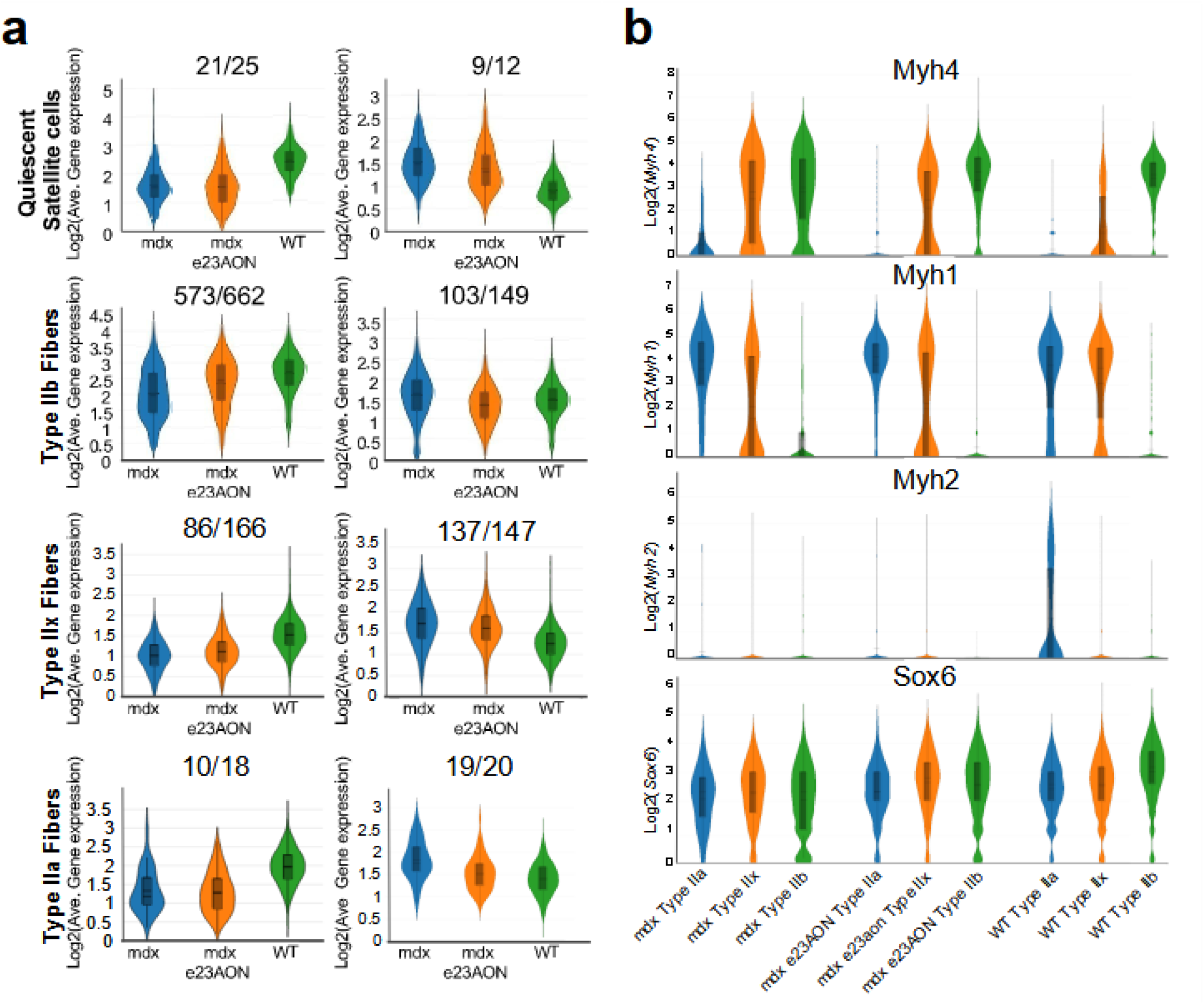
e23AON treatment in *mdx* shifts Myofibers towards more WT behavior (**a**) Average expression of genes upregulated (left) or downregulated (right) in WT relative to untreated *mdx* in quiescent satellite cells (1^st^ row), Type IIb Fibers (2^nd^ row), Type IIx Fibers (3^rd^ row), and Type IIa Fibers (4^th^ row). Y axes = log2 average UMI for differentially expressed genes. The number of DE genes either up (left) or down (right) regulated in WT is given as the denominator above each plot, the numerator indicates how many of these genes are similarly affected by e23AON treatment. Average gene expression changes for additional cell types shown in Figure S5, Gene lists in Data file S2. (**b**) Distributions of expression levels of *Myh4* (1^st^ row), *Myh1* (2^nd^ row), Myh2 (3^rd^ row), *Sox6* (4^th^ row) among Type IIa (blue), Type IIx (orange) and Type IIb (green) fibers of *mdx, mdx* e23AON, and WT mouse muscle. Y-axis = log2 counts of *Myh1, Myh2, Myh4*, and *Sox6* genes for each fiber type/treatment status.

**Figure 6.**
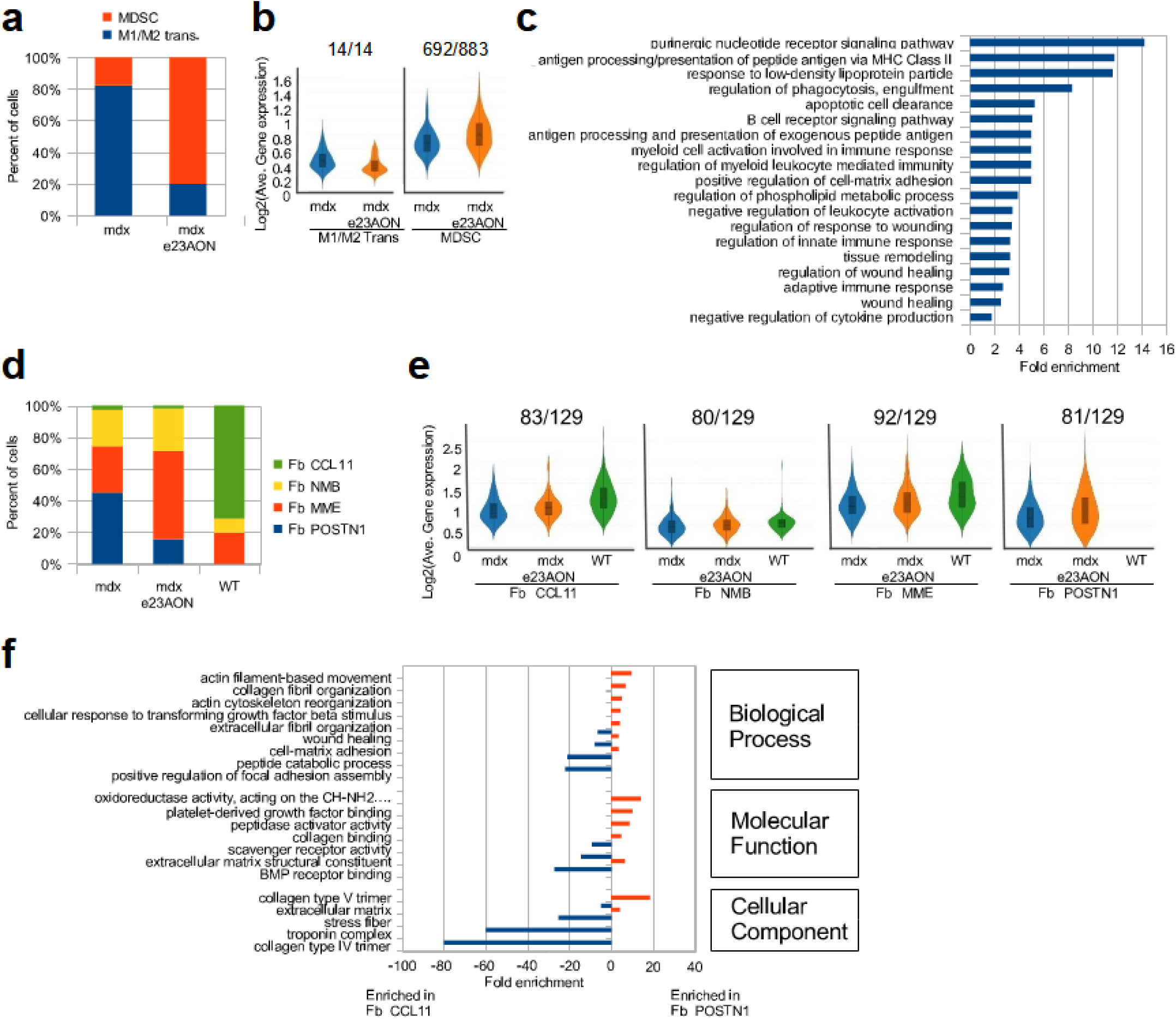
e23AON treatment in *mdx* shifts multiple distinct cell types towards more WT pattern (**a**) Depletion of M1/M2 trans and expansion of MDSC with e23AON treatment. Y-axis shows the relative percentage M1/M2 trans and MDSC populations in *mdx* and *mdx* e23AON. (**b** (**right**)) Log2 average UMI for genes upregulated in MDSC relative to M1/M2 trans (Y-axis), for *mdx* and *mdx* e23AON MDSC populations. Denominator indicates the number of genes upregulated in MDSC, numerator indicates how many of these genes are also upregulated in MDSCs with e23AON treatment. **(left)** Log2 average UMI for genes downregulated in MDSC relative to M1/M2 trans (Y-axis), for *mdx* and *mdx* e23AON M1/M2 trans populations. Denominator indicates the number of genes downregulated in MDSC, numerator indicates how many of these genes are also downregulated in M1/M2 trans with e23AON treatment. (**c**) Representative list of enriched GO terms (Y-axis) based on genes upregulated in MDSC relative to M1/M2 trans. Fold enrichment indicated on x-axis. List of DE Genes between MDSC and M1/M2 transitional cells listed in Data file S3. (**d**) Relative changes in cell proportions of Fb CCL11 and Fb POSTN1 populations and WT intermediates (Fb MME and Fb NMB) between *mdx, mdx* e23AON and WT conditions, Y-axis shows percentages of cell types. (**e**) log2 average UMI of genes upregulated in Fb CCL11 relative to Fb Postn1 (Y-axis) for *mdx, mdx* e23AON, and WT in Fb CCL11 (far-left), Fb NMB (mid-left), Fb MME (mid-right), and Fb POSTN1 (far-right) populations. Above each plot, the denominator indicates the number of genes upregulated in Fb CCL11, and numerator indicates how many of these genes are also upregulated with e23AON treatment. List of DE Genes between Fb POSTN1 and Fb CCL11 cells listed in Data file S4. (**f**) Representative list of enriched GO terms (Y-axis) based on genes differentially expressed between Fb CCL11 and Fb POSTN1. Fold enrichment indicated on x-axis, >0 shows GO terms enriched for genes upregulated in Fb POSTN1, <0 shows GO terms enriched for genes upregulated in Fb CCL11.

In quiescent satellite cells (Fig. 5a, 1^st^ row), 84% (21/25) of genes significantly upregulated in WT relative to *mdx* were also upregulated in response to e23AON. Among these genes, *Pde4b* and *Pde10a* affect the severity of dystrophinopathy, and *Malat1, Kcnma1*, and *Hs6st3* play roles in myogenesis (Table S1b). Further, 75% (9/12) of genes significantly downregulated in WT relative to *mdx* were also downregulated with e23AON treatment, and their average expression in *mdx* e23AON were normalized toward the WT distribution of expression (Fig. 5a, 1^st^ row, medians: *mdx* = 3.32, *mdx* e23AON = 3.00, WT = 2.58). Many of these genes are associated with satellite cell activation (*Gpc6*), asymmetric division (*Egfr*), myoblast differentiation (*Runx1*), muscle growth/regeneration (*Grb10, Zbtb16, Cdon, Notch3, Ror1*), muscle diseases (*Mybpc1, Lama2*), or aging muscle (*Rbms3, Gm42418*) (Table S1b). Overall, the genes upregulated in *mdx* satellite cells relative to WT indicated that they appear primed for activation. The decreased expression of these genes in e23AON-treated *mdx* indicated a partial recovery towards WT physiology.

Gene expression of mature myofiber nuclei in *mdx* were substantially affected by lack of dystrophin, and there was a striking similarity of gene expression perturbation across *mdx* myofiber nuclei type relative to WT: 79% of genes differentially expressed in type IIx fibers were also differentially expressed in type IIb, similarly 95% of genes differentially expressed in IIa are also differentially expressed in type IIb and type IIx, and the majority of these are also differentially expressed in MTJ, Tcap, immature fibers, and partly mature myofiber nuclei (Datafile S3). Most of these differentially expressed genes were also differentially expressed between *mdx* and e23AON-treated myofiber nuclei (68% in type IIx, 41% in type IIa, 78% Type IIb), with a strong trend towards e23AON-treated nuclei recovering a more WT gene expression (Fig. 5a). Many of the differentially expressed genes have been associated with muscular dystrophy (i.e., *Dmd, Lama2*, and *Cd36*) (Datafile S3, Table S1b). While genes associated with myogenesis (*Sox6, Mybpc1, Setbp1*), muscle contraction (*Myh1, Myh2, Myh4*), and muscle strength (*Actg1, Bin1, Dmpk*) were expressed in both *mdx* and WT myofibers, genes associated with improved stress recovery (*Car3, Ucp3, Agbl1, Pdk4*) were only upregulated in myofibers from WT and e23AON-treated *mdx* (Datafile S3, Table S1b).

In *mdx*, myofibers mis-regulate genes required for proper terminal differentiation (*36*). We identified *Sox6*, a gene essential for terminal differentiation of fiber types (*36*), as suppressed in untreated *mdx* IIx fibers (p-value =8E-9, FC = 3.5, Fig. 5b, Datafile S3) and IIb fiber nuclei (p-value = 1E-263, FC = 2.0, Fig. 5b, Datafile S3) whereas *Myh2, Myh1* and *Myh4* markers of Type IIa, Type IIx and Type IIb fiber nuclei respectively, were upregulated in the wrong myofiber types in *mdx* (Data file S3, Table S1b). e23AON treatment partially restored WT expression patterns, reducing expression of *Myh1* in IIb fibers (p-value = 1E-24, FC = 0.55, Fig. 5b, Datafile S3) and *Myh4* in IIx fibers (p-value = 4E-13, FC = 0.77, Fig. 5b, Datafile S3), and increasing the expression of *Sox6* in IIx fibers (p-value = 2E--52, FC = 1.5, Fig. 5b, Datafile S3), and while not statistically significant IIa and IIb also trend in the same direction. These findings suggest that loss of dystrophin results in fiber dysmaturation which can be partially reversed with AON dystrophin rescue.

We observed a substantial shift in the composition of the myeloid compartment in *mdx* versus *mdx* e23AON and WT, with ratios of M1/M2 transitional to MDSC in *mdx* inverting in response to e23AON skipping (Fig. 6a, Table 1). To probe potential consequences of MDSC expansion and concomitant M1/M2 transitional cell contraction in e23AON treated *mdx* muscle, we identified DE genes between these two populations (Datafile S4). This analysis revealed upregulation of genes basic to MDSC biology including phagocytosis, autophagy, exosome function and other processes (*37*) (*38*) (Fig. 6 b, c). Further, regulators of: fatty acid oxidation (FAO) metabolism (*Sat1, Ac49090, Stard5, Ppar, Lacc1, Axl3.4, Ddh, Lair, C1qc, C1qb Cd36, Mertk*); immune suppression (*Trafd1, Cd180, Trpm7 Clec12a, Cd45, Pirb, Tyrobp, Trem2, Fcgr2b, Cd300c, Cd300a, Gpmnb Inpp5d, Il-10r, Tgfbr1, Ccr5, Itgav, Adam17, Il-15,Ccr2*); anti-inflammatory M2 skewing (*Tmsb4x, Hdac8, Trpm7, P2ry6, Ptbp3, Pias, Sirpα, Cd47, Stat5b*) and muscle regeneration (*Igf1, Il15*) were all enriched in MDSC versus transitional M1/M2 cells. Of note, M2MDSC and MDSC populations both expressed genes characteristic of FAO including *Lpl, Pparγ*, and *Cd36* (Datafile S4, Table S1c). Likewise, the M2 population present in WT and expanded in AON treated muscle express genes characteristic of anti-inflammation and wound healing including MERTK, CD36, C1qc and others).

Analysis of differentially expressed genes between MDSC and M1/M2 transitional populations indicated that e23AON treatment resulted in immune cell reprogramming towards a more “MDSC-like” gene profile (Fig. 6b): The average expression of genes upregulated in MDSC relative to M1/M2 transitional cells is further upregulated in MDSCs with e23AON treatment (Fig. 6b, right). Likewise, the average expression of genes downregulated in MDSC relative to M1/M2 transitional cells is further downregulated in M1/M2 transitional cells with e23AON treatment (Fig. 6b, left).

Within fibroblast populations, Fb CCL11 predominated in WT muscle, representing 7.5% of intramuscular nuclei, whereas in *mdx* Fb CCL11 represented only 0.5%. Most notably, in *mdx* muscle, Fb POSTN1, a population absent in WT muscle, expanded to 8.8% of all nuclei along with Fb NMB (4.49% *mdx* vs .93% WT) and Fb MME (5.67% *mdx* vs 2.07% WT). Exon skipping lead to significant contraction of Fb POSTN1 (2.98% *mdx* e23AON vs 8.8 % *mdx*) but doubling of Fb NMB and Fb MME with treatment. Based on lineage analysis (Fig.1d, lower), we interpreted this as a redistribution from Fb POSTN1 towards Fb MME and Fb NMB, that are potential intermediates between Fb POSTN1 and Fb CCL11 (Fig. 6d, Table 1, Datafile S3). Further, there was a more WT (Fb CCL11) pattern of gene expression within Fb MME: 87% (39/45) of genes upregulated in WT relative to *mdx* also had a higher average expression in *mdx* e23AON (Fig. S4A), and 94% (16/17) of genes downregulated in WT relative to *mdx* were also downregulated in e23AON-treated *mdx* relative to *mdx* (Fig. S4A). Genes upregulated in Fb MME include *Lama2* (loss is associated with muscular dystrophy (*39*)), *Plxdc2* (inflammation suppressor (*40*)), and *Cxcl14*, (implicated in chemotaxis (*41*)). Downregulated genes in e23AON-treated *mdx* Fb MME include *Sparc, Meg3* and collagens, *Col3a1, Col1a1*, and *Col1a2*, which have been implicated in inflammation or fibrosis (Datafile S3, Table S1d, Fig. S4A).

Across fibroblast types, there was an overall shift toward WT-predominant Fb CCL11 gene expression pattern in response to e23AON treatment. Of 129 genes significantly upregulated in Fb CCL11 relative to Fb POSTN1, with e23AON treatment: 83 were upregulated in Fb CCL11, 80 in Fb NMB, 92 in Fb MME, and 81 in Fb POSTN1. Even the most pro-dystrophic/fibrotic fibroblasts, were shifted toward a more WT gene expression pattern with dystrophin induction (Fig. 6e). There was significant enrichment of genes upregulated in Fb CCL11 with functions including wound healing, extracellular matrix, and production of collagen fibrils (Fig. 6f), while many genes enriched in Fb POSTN1 (Datafile S5), were pro-fibrotic/dystrophic in skeletal muscle, including, *Mmp3, Tnc, Col8a1, Bcat1, Ltbp2, Wisp1, Postn1* and *Ctgf ((42, 43*) and Table S1d).

To identify fiber type, immune and fibroblast populations using well described protein markers, we stained tissue sections with antibodies directed against: 1) Myh2, Myh1, Myh4 to visualize fiber types IIa, IIx, and IIb, respectively, CD45 to visualize all immune cells and CD206 to specifically visualize M2 macrophages, and 3) POSTN1 to visualize profibrotic fibroblast markers.

Type IIa fibers, which make up 15% of fibers in WT, were nearly absent (3% of fibers) from untreated *mdx* TA. With treatment the representation of type IIa fibers was partly restored to 8%, consistent with snRNAseq inferences (Fig. 7). Visualization of CD45, CD206 and POSTN1 enabled observation of spatial relationships between M2 macrophages and POSTN1 Marked fibrotic lesions. *mdx* TA show densely packed CD45+ immune infiltrates both within and around fibrotic lesions, whereas CD206 high expressing M2 cells are preferentially found surrounding the fibrotic area, (Fig. 8). In *mdx* e23AON treated TA shows CD45+ and CD206+ M2 macrophages distributed around individual myofibers or in smaller POSTN1 marked fibrotic foci. WT muscle shows minimal fibrosis with the predominant immune cells expressing the CD206 M2 marker scattered across the muscle tissue (Fig. 8). These findings are in keeping our findings using snRNAseq and highlight the value of assessing spatial distribution of intramuscular cell subsets in guiding our understanding of dynamic relationships between immunity and fibrosis in the context of dystrophin of and repair.

**Figure 7.**
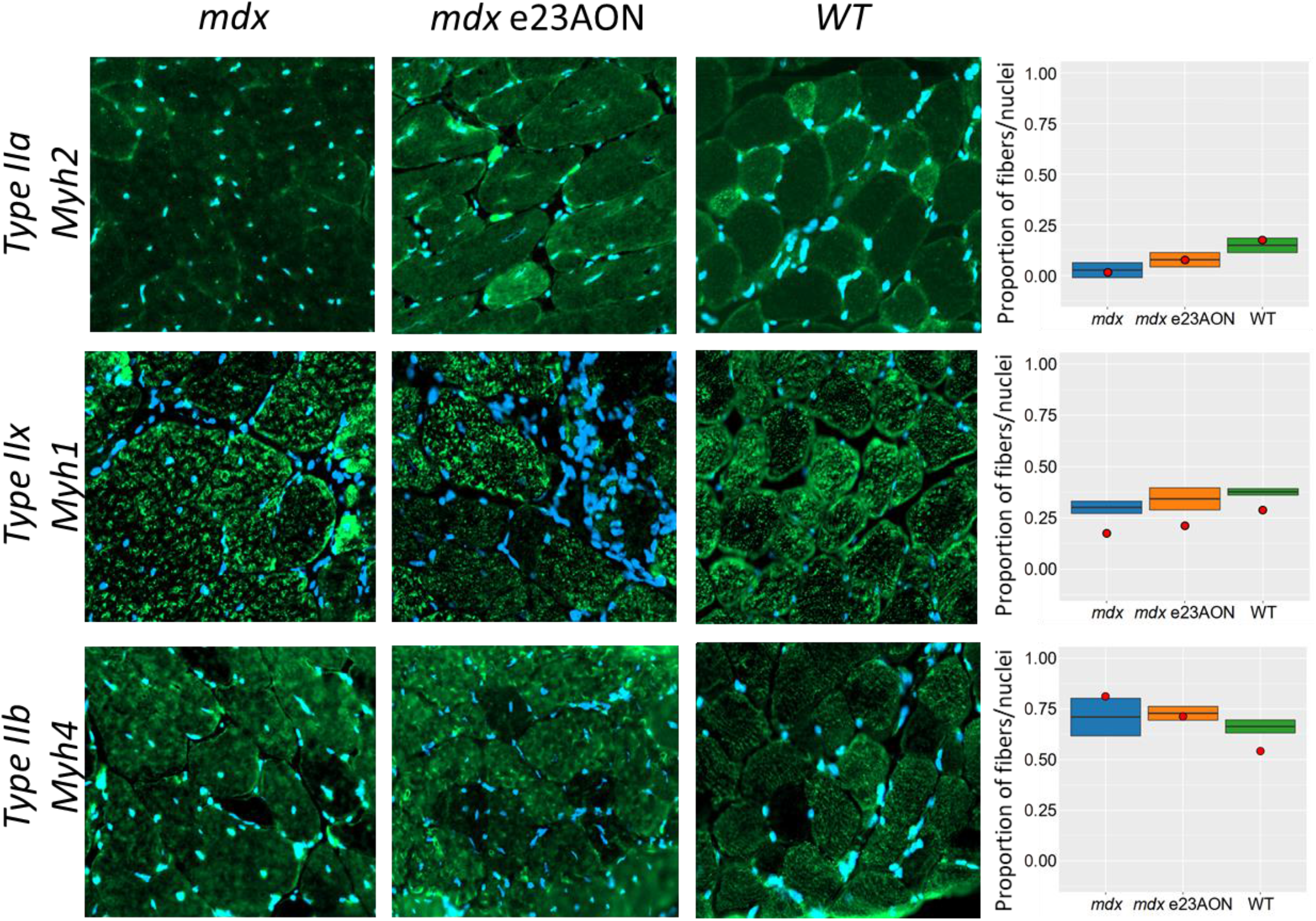
Myosin staining reveals preferential Fiber type degradation and recovery in e23AON treated *mdx*. MYH2 (1^st^ row), MYH1 (2^nd^ row), MYH4 (3^rd^ row) were stained (Green) in muscle sections from *mdx* (left) *mdx* e23AON (center) and WT (right) mice, in combination with DAPI (blue) to mark nuclei. Regions were chosen to best reflect the distribution of fibers across each tissue. Quantification of the proportion of fiber type staining positive for MYH2, MYH1, or MYH4 in representative regions on the muscle across all samples are shown to the right of the IHC images. Thick line shows mean proportion across all samples in each treatment condition, box shows 1 standard deviation. Red dot indicates the proportion myonuclei for each respective fiber type for each treatment condition,

**Figure 8.**
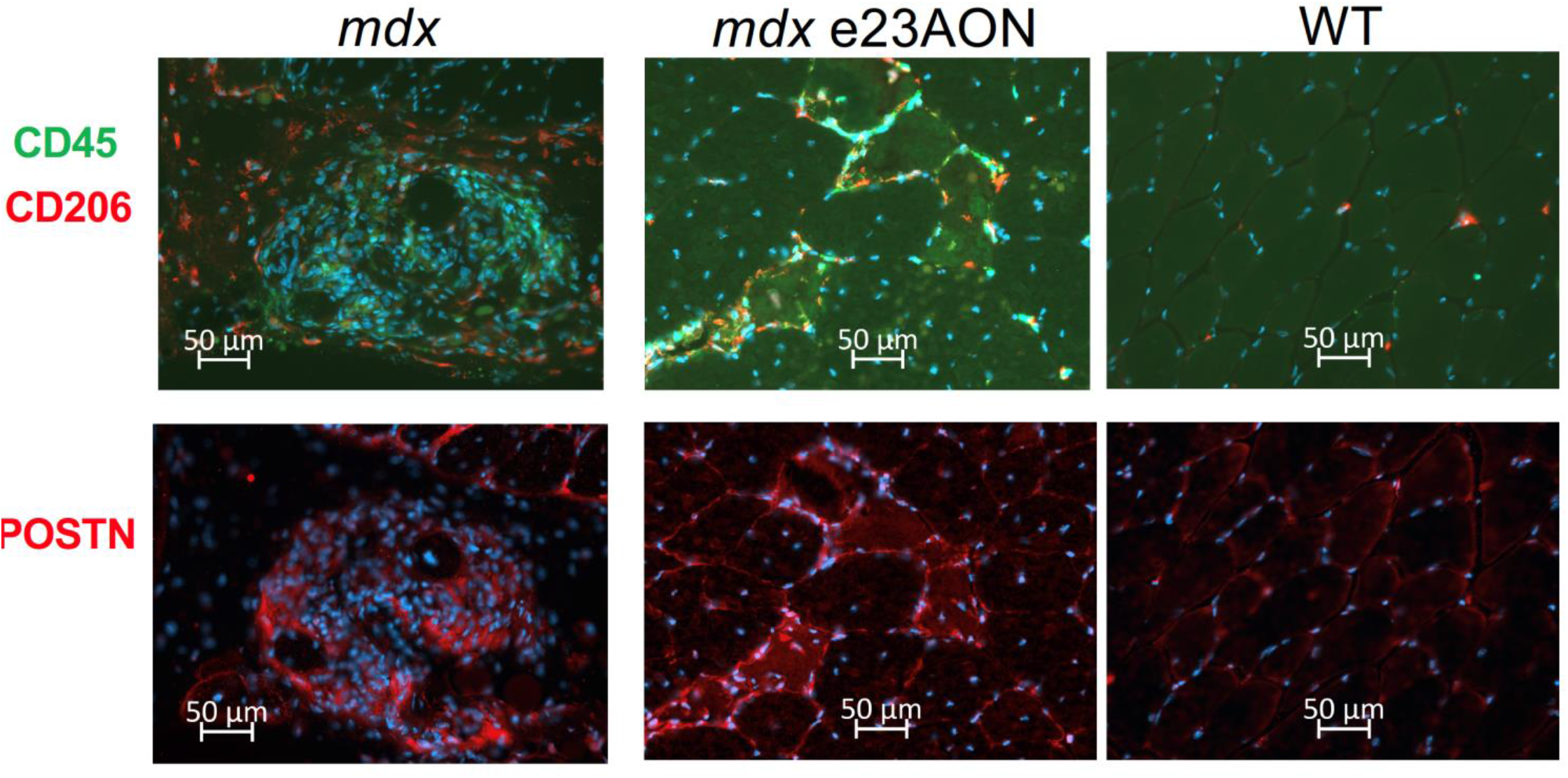
*mdx* and *mdx* e23AON mice show POSTN1 expression within fibrotic lesions surrounded by predominantly peripheral M2 macrophage CD45 (green) and CD206 (red) were stained in muscle sections from *mdx* (top left) *mdx* e23AON (top center) and WT (top right) mice, in combination with DAPI (blue) to mark nuclei. Regions were chosen to best reflect number and type of immune cells present and differences in levels of immune infiltration and POSTN1 expression occurring with disease and treatment. POSTN1 protein was probed in fibrotic lesions in sections from *mdx* (2^nd^ row left), in *mdx* e23AON (2^nd^ row center) and WT (2^nd^ row right) mice

### snRNAseq on frozen biopsy sections identifies diverse cell types and gene expression profiles in healthy and DMD human individuals

We analyzed frozen human muscle biopsies from 3 DMD patients and 2 healthy controls, identifying 28 distinct cell types in VL muscle, broadly comparable to cell types identified in murine samples (Fig. 9). Like the comparison of *mdx* and WT, type IIa and IIx fibers and smooth muscle declined in DMD muscle compared to healthy controls, whereas satellite cells, myeloid, fibroblasts, T cells, B cells, and endothelial cells increased in DMD muscle. Both human and mouse showed an increase in adipocytes accompanying loss of dystrophin, though human dystrophic tissue had a greater proportion of adipocytes, consistent with prior reports (*44*). Upon sub-clustering of all samples, we identified 6 discrete myofiber nuclei types, 4 smooth muscle cell types, 4 types of macrophages, 3 sub-types of endothelial, and 5 types of fibroblast, two of which resemble previously reported sub-types, marked by *PRG4* and *LUM* (*7*). In DMD subjects, fibroblast clusters Fb 1, 2, and 4 were expanded and Fb 1 and Fb 2 expressed high levels of *CXCL12*, which encodes a chemokine known to recruit macrophages and T cells to muscle and is required for muscle regeneration (*45*) (*46*). In addition, we confirm previously reported expression of CXCL12 in DMD endothelial cells relative to healthy control(*47*).

**Figure 9.**
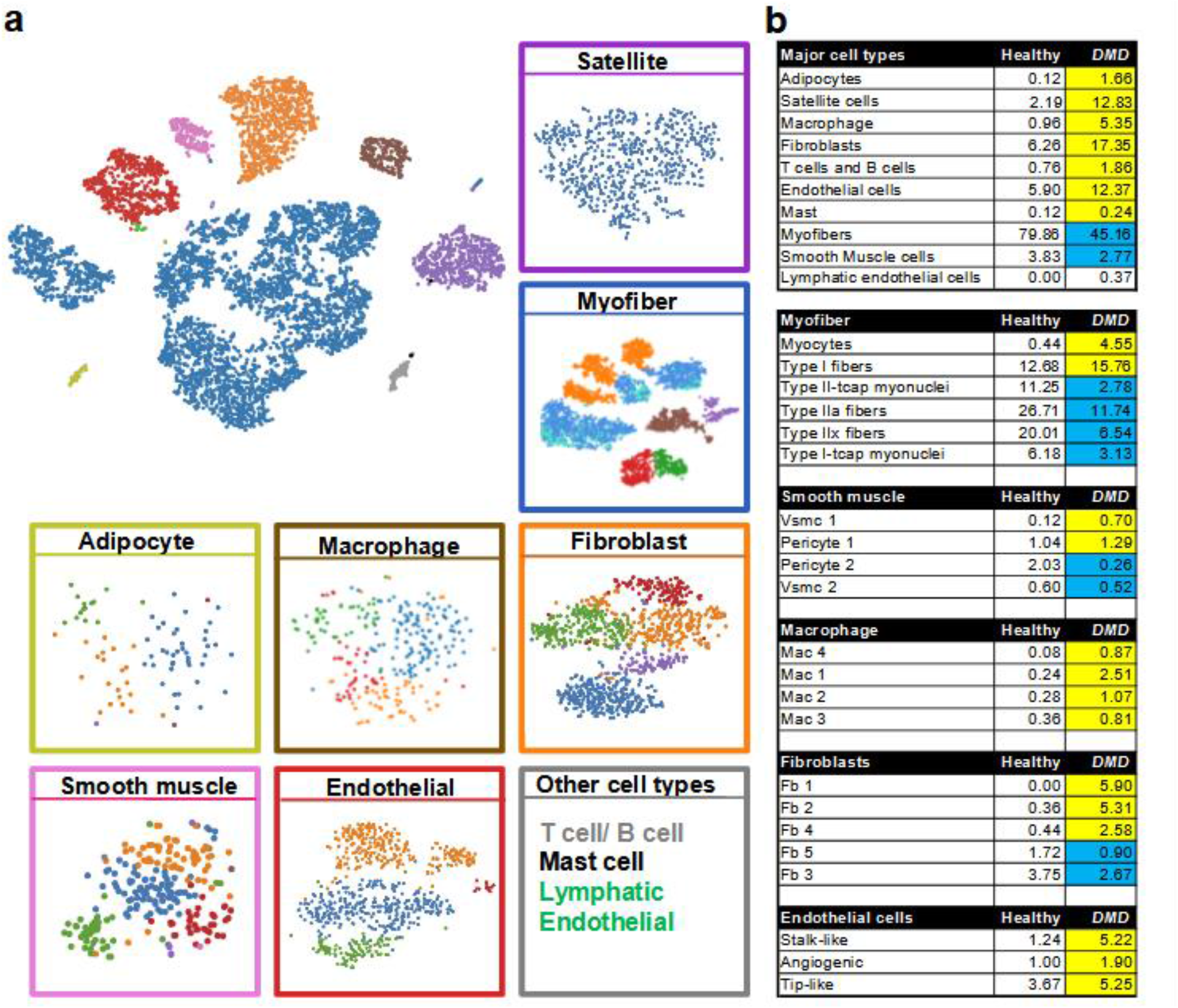
Identification of diverse cell populations in human vastus lateralis from healthy and DMD individuals using snRNAseq. (**a**) TSNE clustering of human VL single nuclei transcripts. Major cell types were identified (top left panel) and re-clustered separately to investigate substructure within each group (box colors surrounding subcluster correspond to cell colors in top left panel). Cell cluster types too small to be easily seen are listed in “Other cell types” panel, text color reflects the color of the population reflected in the top left panel. (**b**) Frequencies of cell populations identified in healthy and DMD muscle. Percentages for each cell population named in the 1^st^ column for healthy individuals and in the 2nd column for DMD affected individuals. Following the major cell types, cell types are further broken down to their respective subclusters. Yellow = increase and Blue = decrease in DMD percentage relative to healthy. Subsets within categories are listed in order of decreasing magnitude of difference between DMD and healthy. Gene names are listed in Data file S5, and Figure S5. Genes differentially expressed between healthy and DMD are listed in Data file S6.

Within the human macrophage/myeloid immune compartment, we identified putative M1, M2, pDC and MDSC-like subpopulations. These populations have the mixed, labile quality previously reported for *in situ* tissue macrophage, in that they express markers consistent with multiple canonical cell types (*48*). Mac 1 was most consistent with an M2 macrophage, expressing CD14, modest levels of *FcγrIIIa/Cd16*, as well as M2 markers *CD23/FCER2, CCL18, CD209* and *F13A1 (48) (*statistically significantly upregulated relative to other clusters), though we also identified markers more typical of an inflammatory state, such as *IL-2RA (49), (50*). Mac 2 most resembled M1 macrophages, expressing *CD14*, and M1 markers; *CD44 (osteopontin receptor), CD86, TLR2 (51), IL-15 (51) (48*), and highest levels of *FcγrIIIa/Cd16* (*48*), Mac 3 is potentially a pro-inflammatory pDC population, most clearly marked by significantly upregulated pDC marker *HFM1* and *FAAH (52*) and Human protein atlas (https://www.proteinatlas.org/)). Mac 4 resembled MDSC or other immune suppressive myeloid cell types in that it was the major population to express *GPNMB*/DC-HIL, which encodes a protein characteristic of MDSC and involved in T cell repression (*53*), and S100A4, critical for MDSC survival (*53*). This same cluster, in the DMD samples only, expressed other immune modulatory and/or pro-fibrotic genes, such as *SPP1*, expressed exclusively in Mac 4 (*54*), (*55*) and *TGFB (56*) as well as pro-inflammatory genes, including *CD64/FCGRA* and *CD64b/FCGR1b (57) (48*). Both MDSC and the M1 populations were increased in DMD muscle relative to healthy control muscle (10 fold and 3-fold respectively), highlighting commonalities between human and mouse dystrophy.

## Discussion

We developed and implemented a method to purify individual nuclei from approximately 3mg frozen mouse or human skeletal muscle for single nuclei sequencing. Because some cells are multinucleated (myofibers) while others (immune, fibroblast, satellite) are mononuclear, we implemented protocols for unbiased sampling from all nuclei types resident in muscle to reveal the complex architecture of skeletal muscle in health and disease. This is a powerful means to: 1) reveal the heterogeneity of cells that compose skeletal muscle, 2) identify novel cell types and myofiber nuclei of specialized function, and 3) observe and quantify substantial shifts in intra-muscular cell types and gene expression. We identified 8 mouse and 5 human subpopulations of myofiber nuclei some reflective of specialized nuclei functions, including NMJ, MTG, Tcap, and provide comparative data on immature, partially mature, type I, IIa, IIx and IIb myofibers and 4 smooth muscle subpopulations. Because type I fibers were not observed in mouse TA, and type IIb were not observed in human VL, we do not attempt to make comparisons between mouse and human.

In both mouse and human muscle, lack of dystrophin is associated with chronic myofiber damage, and we show a common depletion of type IIa and IIx myofibers in mouse TA and human VL, with similar overall patterns of activation, expansion and diversification of satellite cells, myoblasts, immune cells, fibroblasts and other populations (*58*). While we were able to identify an increase in rare exon skipping events induced by e23AON in single cells, the frequency of observation was too low to compare skipped versus unskipped myonuclei. It is unclear whether effects of dystrophin rescue on satellite cell dynamics reflect rescue of dystrophin within satellite cells, where it is proposed to control asymmetric division, or due to decreased activation secondary to myofiber rescue. Our lack of observed exon skipping in satellite cells may reflect a relatively low frequency, the relative scarcity of skipped message or limits to our detection. Thus, we cannot exclude satellite cells as a skipping target. Technical improvements to capture and sequence full length *DMD* will improve the utility of snRNAseq in DMD muscle to address this question at single cell resolution.

In contrast to most immune cells which are reduced in treated vs untreated *mdx*, a select myeloid subset, termed MDSC, is substantially expanded in e23AON-treated *mdx* muscle. The most common immune cell type found in WT muscle, M2, also increases two-fold in treated mdx mice. M1/M2 transitional cells, the primary *mdx* immune population observed, contracts with treatment and takes on a more “MDSC like” gene expression profile. Given the relatedness of these populations, we propose that exon skipping treatment my drive M1/M2 transitional cells to become MDSC ND/OR M2 cells.

While initial pro-inflammatory response to muscle damage is essential for satellite cell activation and myogenesis, a switch to immune mechanisms of anti-inflammation and wound healing guide completion of muscle repair (*59*). By sensing fatty acids, PPARγ can trigger a metabolic shift within immune populations, resulting in a switch from glycolytic to OXPHOS metabolism for energy production and select differentiation and activation of M2 macrophages, MDSC and other immune cells associated with immune suppressive, anti-inflammation and tissue repair (*60*) (*61*). Likewise, iC1q and LAIR-1 upregulation can promote the switch from pro-to anti-inflammation by recruiting MERTK dependent production of specialized pro-resolving mediators and IL10 (*62*). Multiple members of these cascades are upregulated in intramuscular MDSC and M2 cells and are increased further with exon skipping and dystrophin rescue, supporting their potential role in facilitating a metabolic shift toward a more healing and anti-inflammatory muscle microenvironment and identifying potential molecular targets for disease modulation.

This initial description of intramuscular cell populations within human DMD and *mdx* mouse muscle provides a framework for a more complete analysis across many dystrophic samples. Deeper characterization of human skeletal muscles with variable disease course, or with longitudinal sampling in response to therapeutic strategies, will more fully describe the cell types and variations in specialized nuclei types within multinucleated myofibers. Our ability to identify cell composition and gene expression profiles in nuclei prepared from small (3 mg) samples of frozen human muscle now makes feasible snRNASeq analysis on frozen human remnant muscle tissue obtained in the context of dystrophin replacement or other DMD therapies, natural history studies, or from small core needle biopsies of unusual cases, enabling exploration of therapeutic and naturally occurring mechanisms of genetic repair and pathogenesis in DMD and other muscle conditions.

## Materials and Methods

### Mouse study design

Eleven mice were selected from a published *mdx* mouse experiment in which replicate mice were treated weekly with antisense morpholino e23AON over 6 months to induce exon skipping and dystrophin expression in skeletal muscle (*4*). Archived frozen tibialis anterior muscle, stored in LN2 from four e23AON-treated *mdx*, four age-matched untreated *mdx* and three age-matched untreated control C57BL/10ScSnJ mice (WT) were compared (*4*). All animal handling was performed in accordance with applicable national and institution specific guidelines: UCLA protocol ARC # 2011-021-21. Human muscle was obtained by needle biopsy and stored as previously reported (*63*).

### Microscopy and Immunochemistry

Dystrophin and spectrin staining were performed as previously described (*4*). Trypan blue images of nuclei and size beads obtained using a Zeiss Axiovert 40 CFL, scaled as indicated in Fig. 1a. For immunofluorescence detection of immune cells: Crosssections (10um) on glass slides were fixed in 4% paraformaldehyde, solubilized with 0.5% triton, blocked with 3% BSA for 1hour, prior to application of primary antibodies. Staining for Myh2, Myh1, and Myh4 were blocked for 5 hours prior to applying primary antibodies. Antibodies used were as follows: anti-CD45; rat, 30-F11 (ebiosicences), CD206; goat poly clonal (R&D), POSTN1; Rabbit poly clonal,(Thermo Fisher), MYH2; 8F72C8 (Sigma-Aldrich), MYH1; 6F12H3 (Sigma-Aldrich), MYH 4; 2G72F10 (Sigma-Aldrich), Rabbit anti-goat Alexa546 polyclonal (Invitrogen), donkey anti-rat Alexa 488 (Invitrogen), and Donkey anti-rabbit dyelight 550 (Thermo Fisher). Primary antibodies were incubated overnight, washed with 0.1% triton 4 times prior to 1 hour incubation with secondary antibodies, and sections mounted with vectashield DAPI mount.

### Nuclei extraction from fresh frozen tissue

Frozen muscle was cryosectioned into 4 to 12 40μm thick cross sections into a 1.5ml tube on dry ice to provide a total of 3mg of tissue. 0.5ml of an ice-cold solution of 0.2μm filtered 1% BSA in PBS with 100U/ml of type IV collagenase and 0.5U/μl RNAse inhibitor (Sigma Ref # 03335402001, Mannheim, Germany) was pipetted onto the sections, and sections were allowed to settle to the bottom. Seven strokes with “A” gap dounce, followed by 7 with “B” gap were performed. The mixture was subsequently filtered through a 70μm filter, and the effluent centrifuged at 600g for 6 minutes at 4C. Pellets were resuspended in 0.5ml of 1% BSA in PBS with RNAse inhibitor (0.5U/μl) and DAPI (10μg/ml), incubated on ice for 60 minutes, then sorted.

### Nuclei sorting

Nuclei were sorted using a BD FACS ARIA II sorter with UV laser and 70μm nozzle. Nuclei were gated for aggregate removal by size using the forward and side scatter channels, and for doublets using both FSC-A vs FSC-H and SSC-A vs. SSC-H. Nuclei falling within the DAPI bright gate were collected into 0.5ml of 1%BSA in PBS with RNAse inhibitor (0.5U/μl). Yields ranged from 6,000 to 60,000 nuclei per sample. Following sorting, nuclei were centrifuged at 600g for 6 minutes and the pellet was resuspended and passed through a sterile 40μm filter (Fisher # 08-771-23, Tewksbury, Massachusetts). Up to 20,000 nuclei, but less than 40μl, was loaded for 10x Chromium library preparation.

### Library sequencing

Within our experiments, optimal sequence data generation resulted from loading 10,000-20,000 nuclei for 10x Chromium Single cell 3’ v3 library construction. All sequencing was performed on an Illumina Novaseq 6000 S2 2×50, according to manufacturer guidelines (10x Genomics, Doc #CG000204). Target read depth was 40,000 reads per nuclei library. Post sequencing, library quality was assessed by Cellranger (10X Genomics) count output.

### Statistical analysis

Raw sequence data for all 11 mice and 5 human samples were processed using Cellranger 3.0.2 (https://github.com/10XGenomics/cellranger) Mkfastq (formation of fastq files) and Count to generate gene expression matrices aligned to custom per-mRNA mm10 (mouse samples) or hg19 (human samples) reference genomes. Doublets, identified using DoubletFinder_V3 (*21*), along with any nuclei <200umi were removed along with mitochondrial genes before aggregating datasets. Major cell type populations were identified through T-SNE dimensional reduction along with k-means clustering (K= 10). Each of these populations were reanalyzed using Cellranger to identify specific cell types.

Statistical analysis of differential gene expression was performed using Seurat 3.2.3 (https://satijalab.org/seurat/) run on R version 3.3.2. Single nuclei data were log normalized and scaled. Within each sub-cluster, marker genes were identified for each cell type using DESeq2 test using FindAllMarkers om Seurat. Significant marker genes were identified with p-value < 0.05 after bonferroni correction. For each identified cell type, we also compared the *mdx* vs WT nuclei, as well as *mdx* vs e23AON-treated *mdx* by gene expression using DESeq2 test using FindAllMarkers. Differentially expressed genes needed p-value < 0.05 after bonferroni correction. Gene ontology enrichment analyses were performed using Ease David passing FDR corrected p-value < 0.05. To determine the relatedness of highly similar cell types we used the Pseudotiming analysis R package Slingshot 1.7.2 (*27*). Cell types that were removed from the main body of the sub-cluster were removed from the analysis. The remaining clusters underwent UMAP clustering, (Seurat Run UMAP based on the Seurat FindVariableFeatures) for use in Slingshot analysis. The R package Harmony was used to correct data to ensure distinct cell types were not the product of batch effects(*64*).

Kmers of length 25 were extracted from mouse *Dmd* sequence NM_007868.6 matching reads that aligned to junctions e22-e23, e23-e24 and e22-e24. FASTQ reads where then pattern matched against the kmers using the linux grep utility. Any read matching a kmer of interest was retained and counted. 10x barcodes were used to correct for PCR duplicates and unique reads were counted.

### Human subjects

All subjects were consented in compliance with institutional and federal laws, and UCLA IRB approvals 18-001366 or 19-00090.

## Supporting information

Datafile S1 - Seurat findallmarkers files for mouse data

Datafile S2 - M2MDSC_vs_MDSC differentially expressed genes

Datafile S3 - Differentially Expressed genes mdx vs mdx e23AON vs WT mouse data

Datafile S4 - MDSC vs. M1.M2 transitional differentially expressed genes

Datafile S5 - FB POSTN1 vs. FB CCL11 differentially expressed genes

Datafile S6 - Seurat findallmarkers files for human data

Datafile S7 - Differentially expressed genes dmd vs control human data

Figure S2: heat maps mouse: A-z of all heatmaps not shown in main text

Figure S5: heat maps human: A-z of all heat maps

Supplemental Figures and References

Table S1: Key genes and references

## Acknowledgments

General: We thank Dr. Jeremy D. Woods, MD, for assistance in collecting muscle biopsies, Dr. Xinmin Li, and technical support from the UCLA Technology Center for Genomics & Bioinformatics (TCGB) core, Iris A. Williams and Dr. Zoran Galić at the UCLA Jonsson Comprehensive Cancer Center Flow Cytometry Core Facility (NIH P30 CA016042).

## Author contributions

MCM and SFN conceived of the project and supervised all analysis and writing.

DDSA, KNC, MCM and SFN wrote the manuscript.

FB, SRN and SFN performed and preserved human muscle biopsies.

EDD consented and scheduled human subjects.

FB and KNC performed all muscle sectioning.

DDSA, KNC, and JW developed and implemented nuclei purification techniques.

DDSA developed and implemented nuclei sorting.

RW devised methodology for identifying skipped *Dmd* RNA in snRNAseq data.

KNC and DDSA performed TNSE and subsequent analysis and identification and quantification of cell population and gene expression signatures.

KNC performed all other analysis for the manuscript data analytical pipeline

DWW, EIM and LR analyzed dystrophin expression included in Figures 3b, Fig S3,

DDSA, KNC, SFN, FB, SNR, RTW, JW, SFN and MCM aided in preparing and reviewing manuscript.

## Data and materials availability

All data associated with this study are available in the main text or the supplementary materials. Accession numbers for public data base will be provided for any data relating to the paper.

## Supplementary Materials

Supplemental references: 65 - 248

Figure S1: Lineage tracing analysis of subpopulations identifies functional relatedness of endothelial and smooth muscle sub populations

Figure S2: heat maps mouse: A-z of all heatmaps not shown in main text

Figure S3: Dystrophin staining of whole murine quadricep from treated and untreated *mdx* mice

Figure S4: e23AON treatment in *mdx* shifts multiple distinct cell types towards more WT behavior (additional cell types)

Figure S5: heat maps human: A-z of all heat maps

Table S1: Key genes and references

Data file S1: Seurat findallmarkers files for mouse data

Data file S2: MDSC vs. M2MDSC differentially expressed genes

Data file S3: Differentially Expressed genes *mdx* vs *mdx* e23AON vs WT mouse data

Data file S4: MDSC vs. M1/M2 transitional differentially expressed genes

Data file S5: Fb POSTN1 vs. Fb CCL11 transitional differentially expressed genes

Data file S6: Seurat findallmarkers files for human data

Data file S7: Differentially expressed human genes; DMD vs healthy control data

