## Supplementary material for "Single nuclei transcriptomics of muscle reveals intra-muscular cell dynamics linked to dystrophin loss and rescue": Figure S2: heat maps mouse: A-z of all heatmaps not shown in main text

a. Myofiber subcluster heatmap

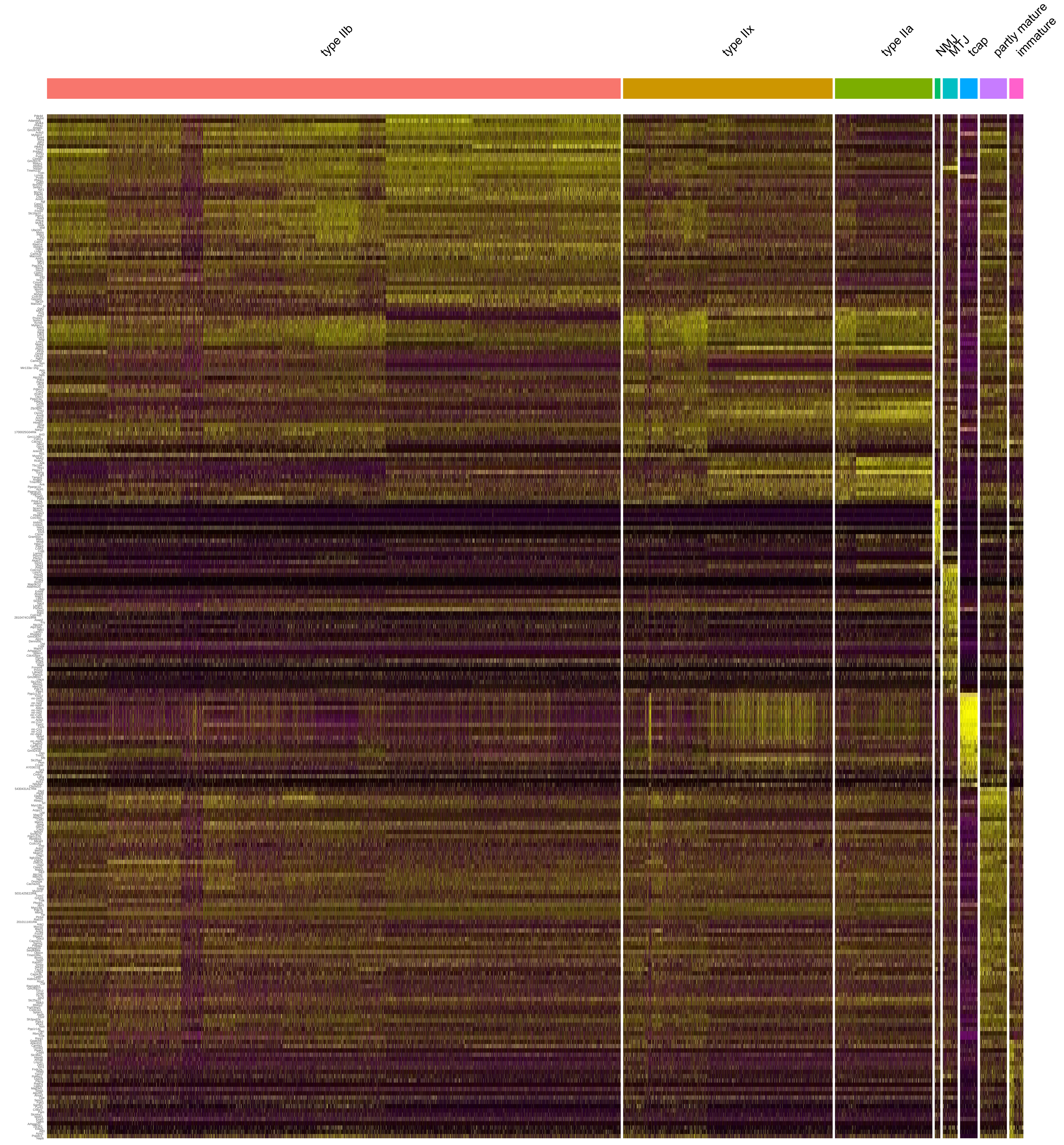


b. Myofiber progenitor (satelite cells/myoblasts) subcluster heatmap

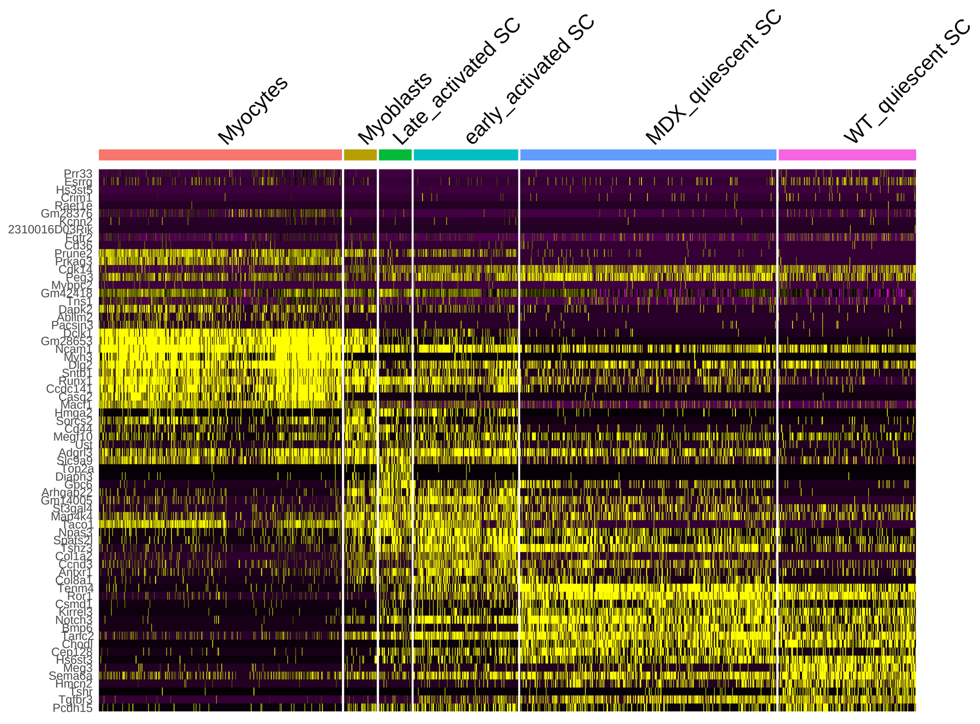

c. Immune cell subcluster heatmap

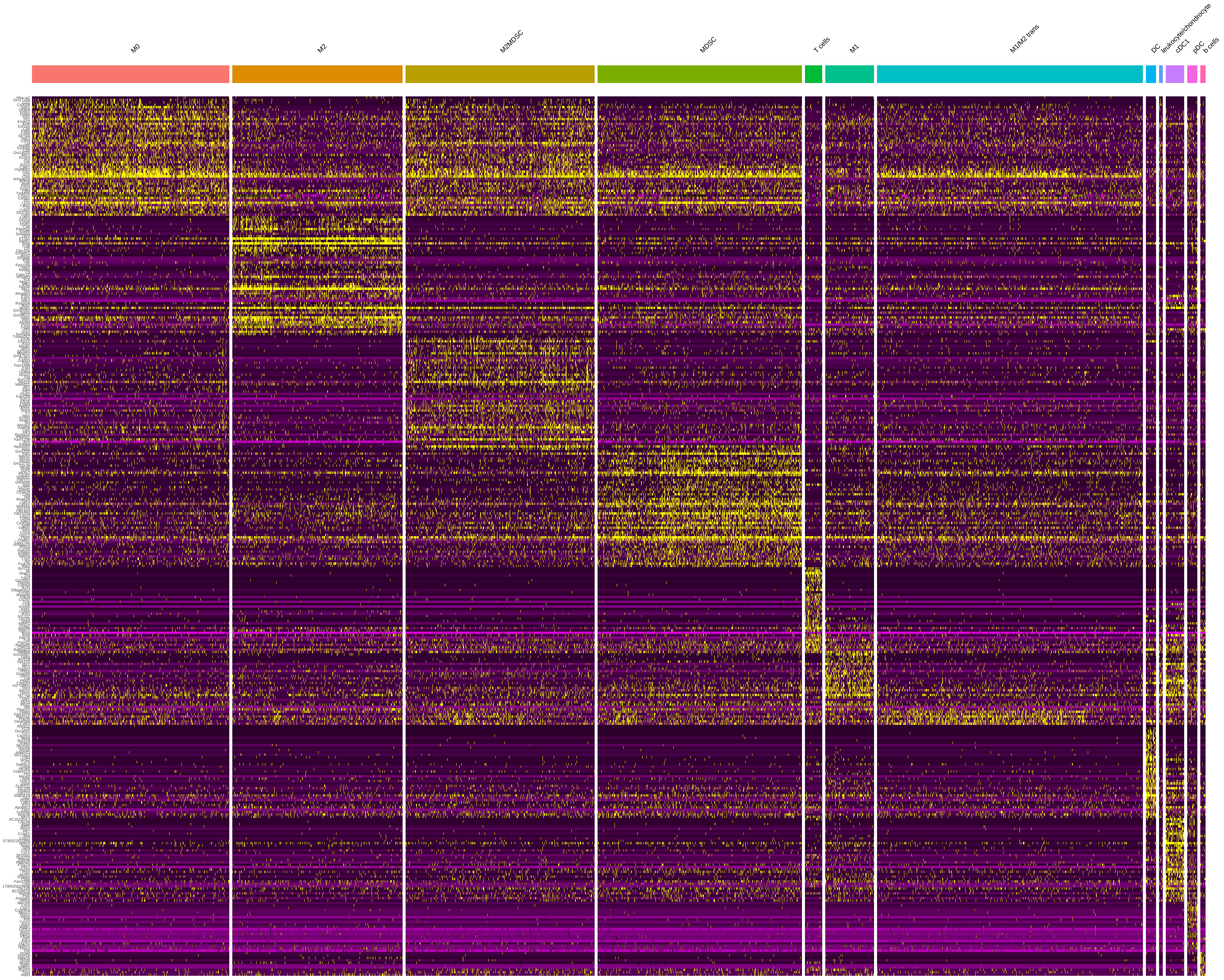

d. Fibroblast subcluster heatmap

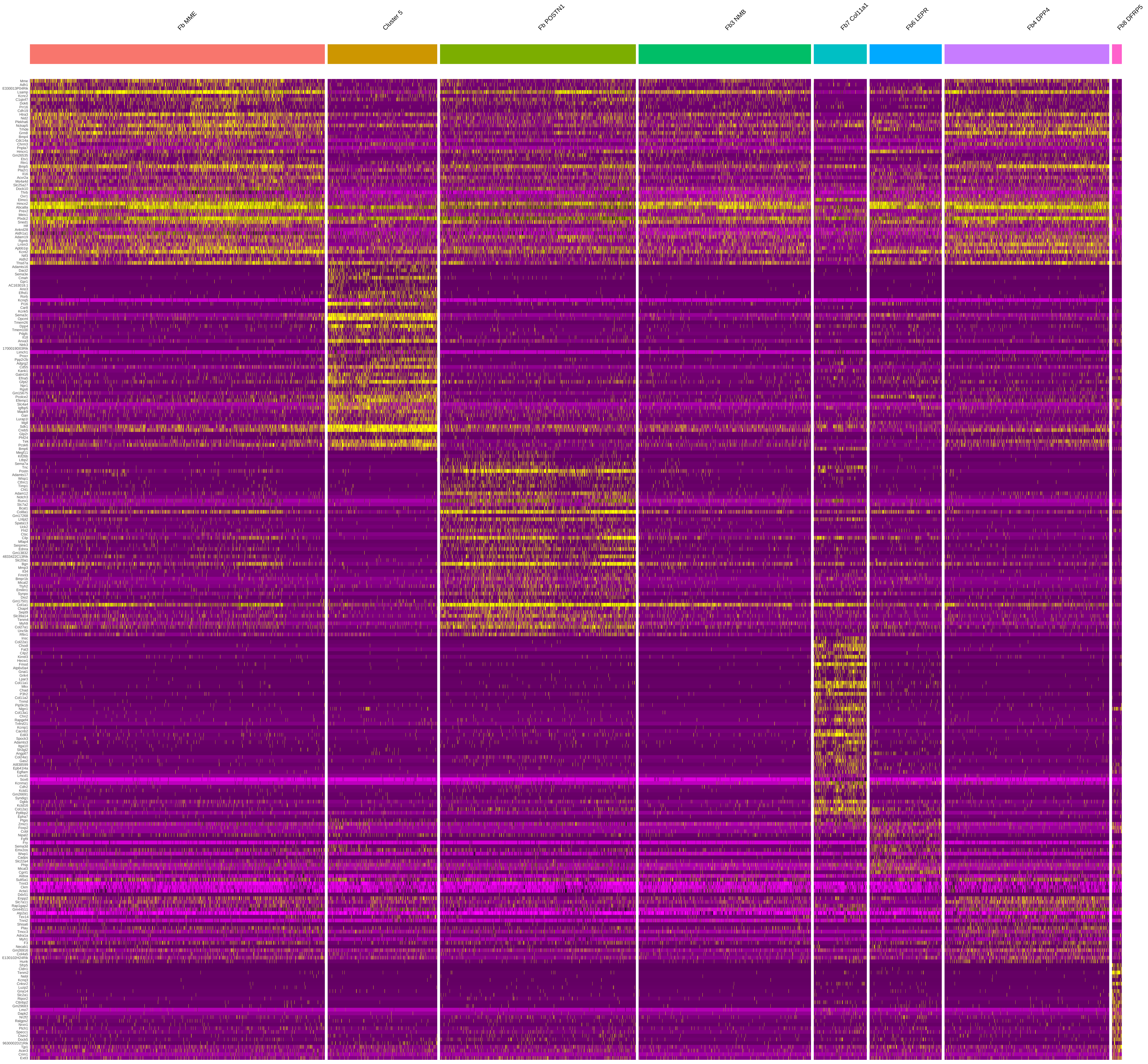

e. Smooth muscle subcluster heatmap

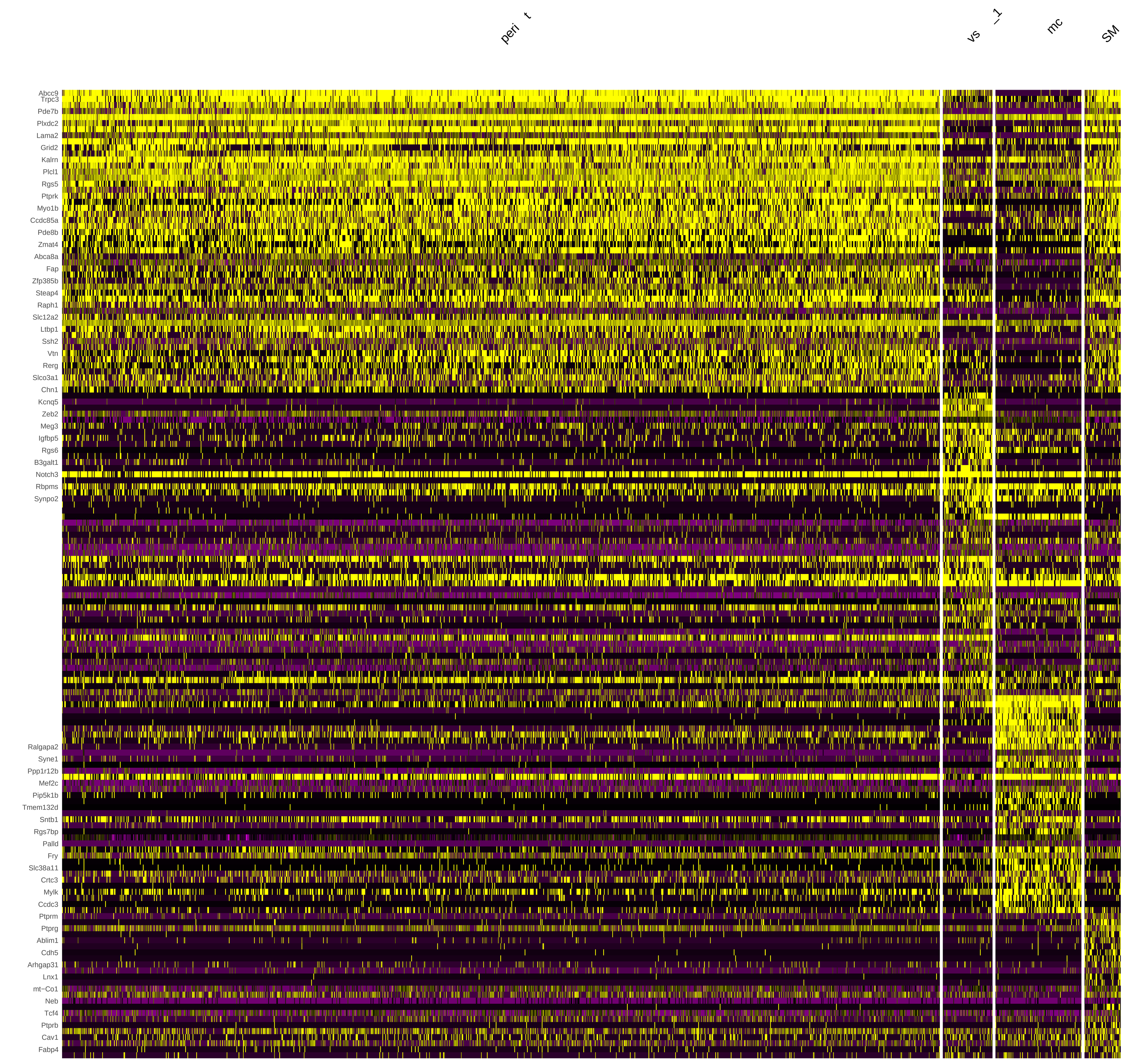


f. Endothelial subcluster heatmap

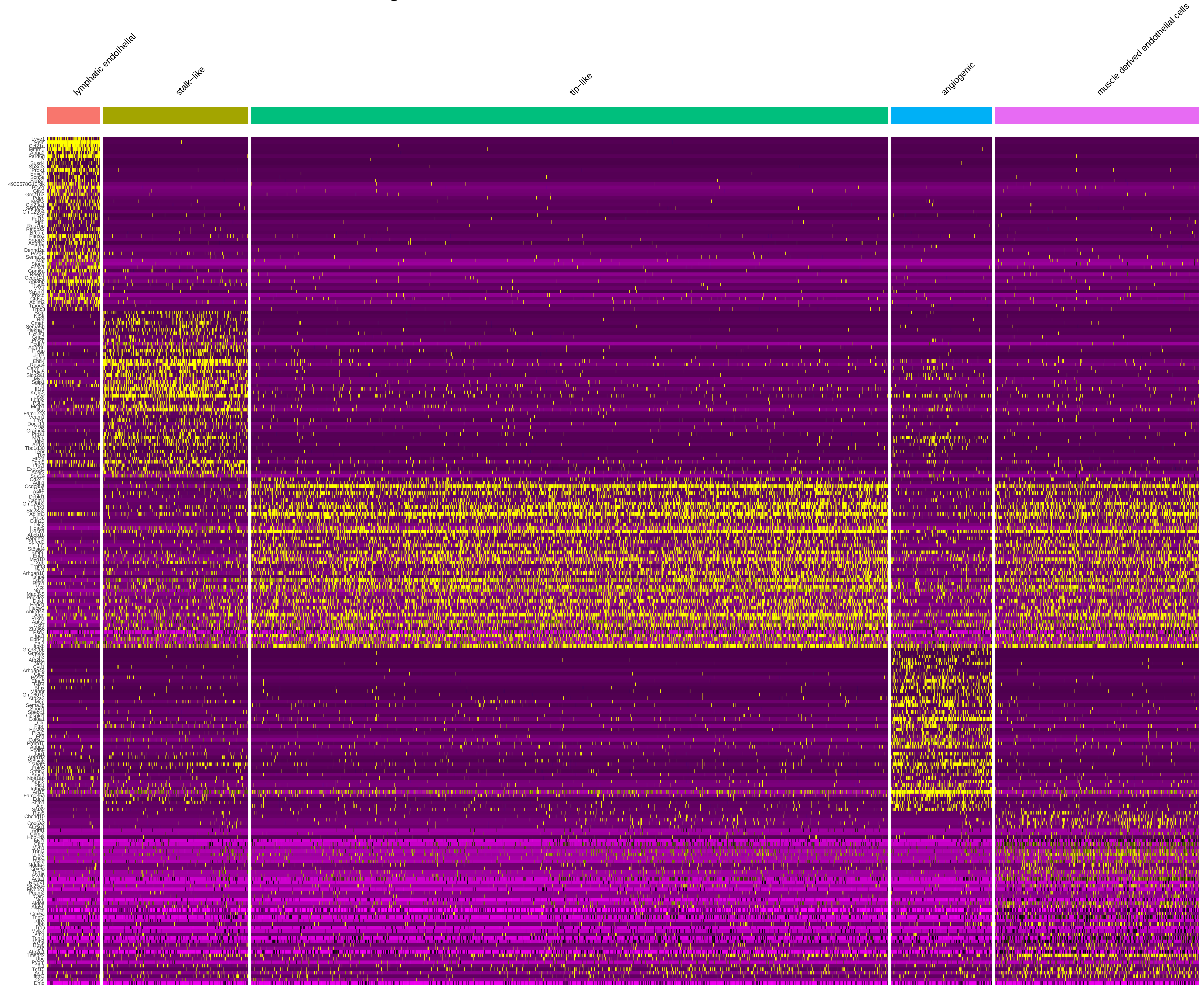

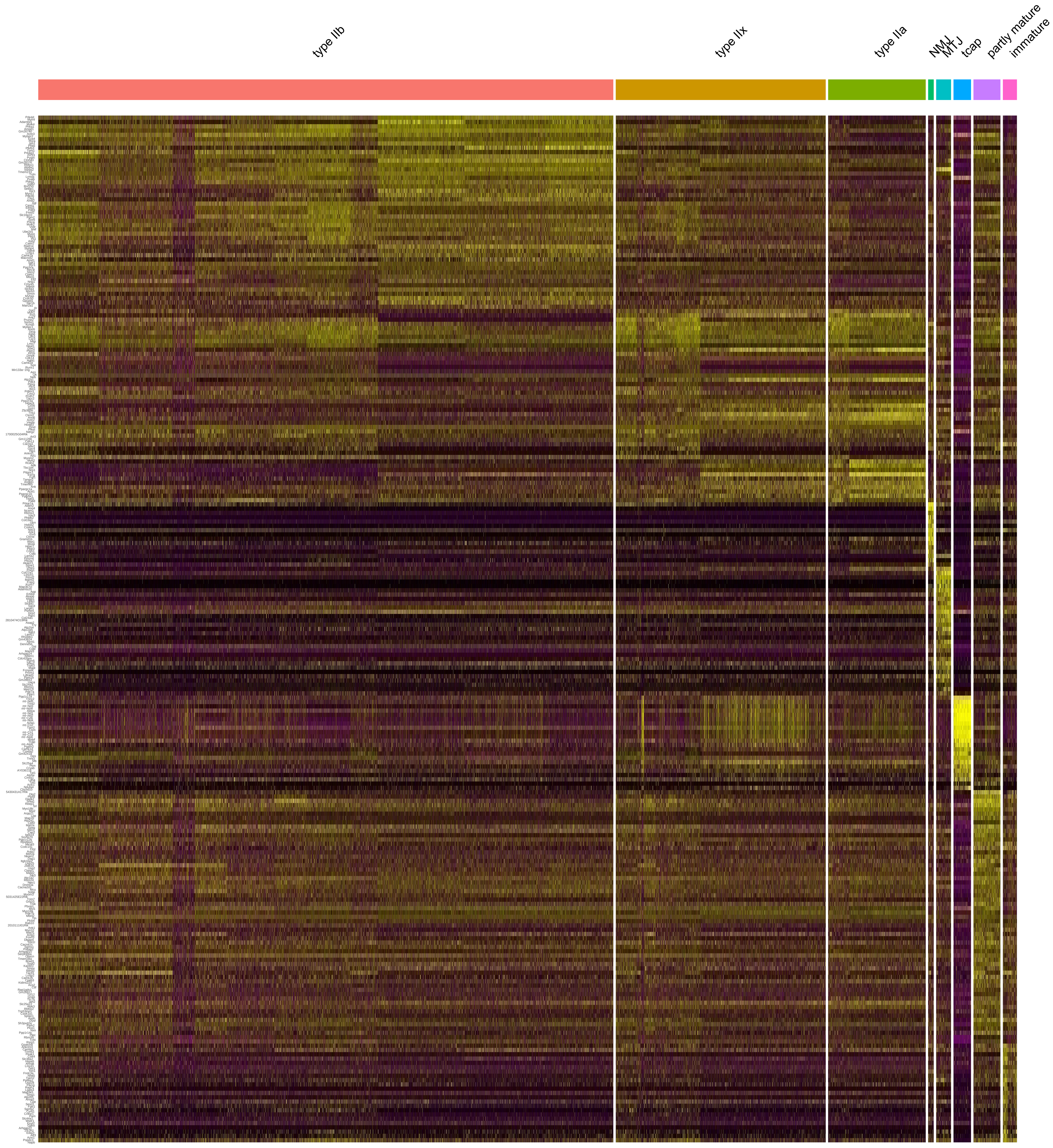
type IIb
type IIx
type IIa
NMJ
MTJ
tcap
partly mature
immature
