## Supplementary material for "Single nuclei transcriptomics of muscle reveals intra-muscular cell dynamics linked to dystrophin loss and rescue": Figure S5: heat maps human: A-z of all heat maps

### a. Human Myofiber heatmap

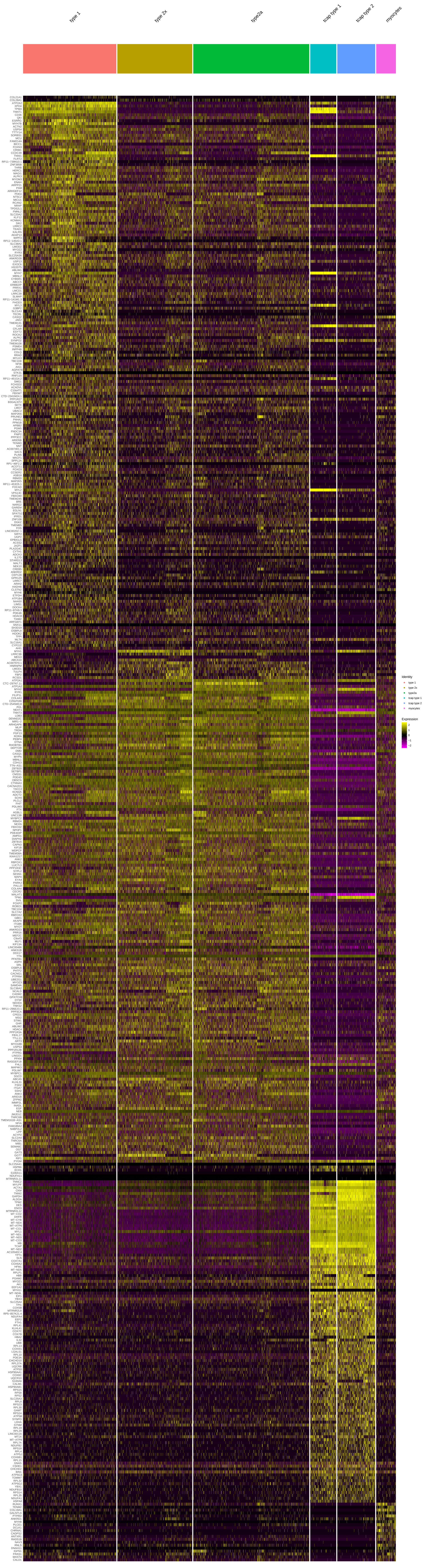

### b. Human Fibroblast heatmap

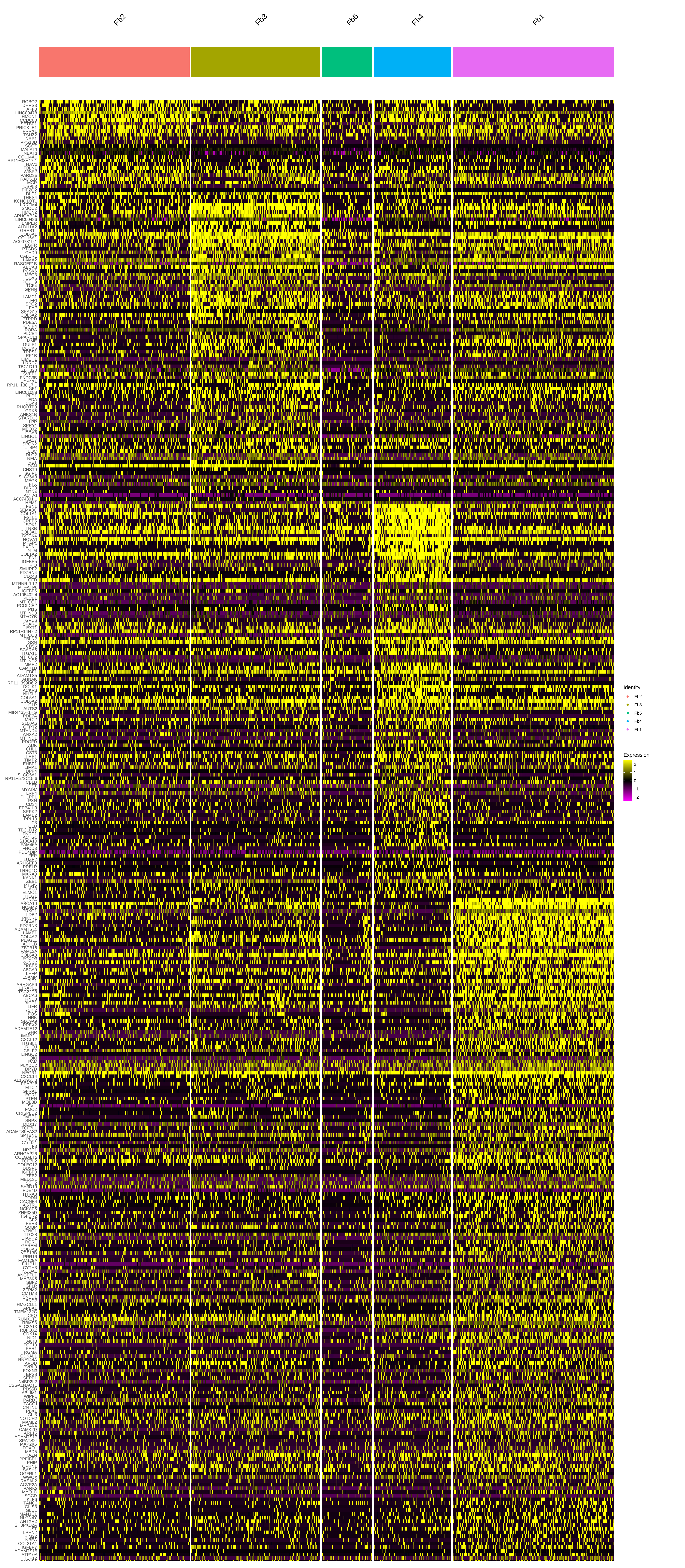


### c. Human Endothelial heatmap

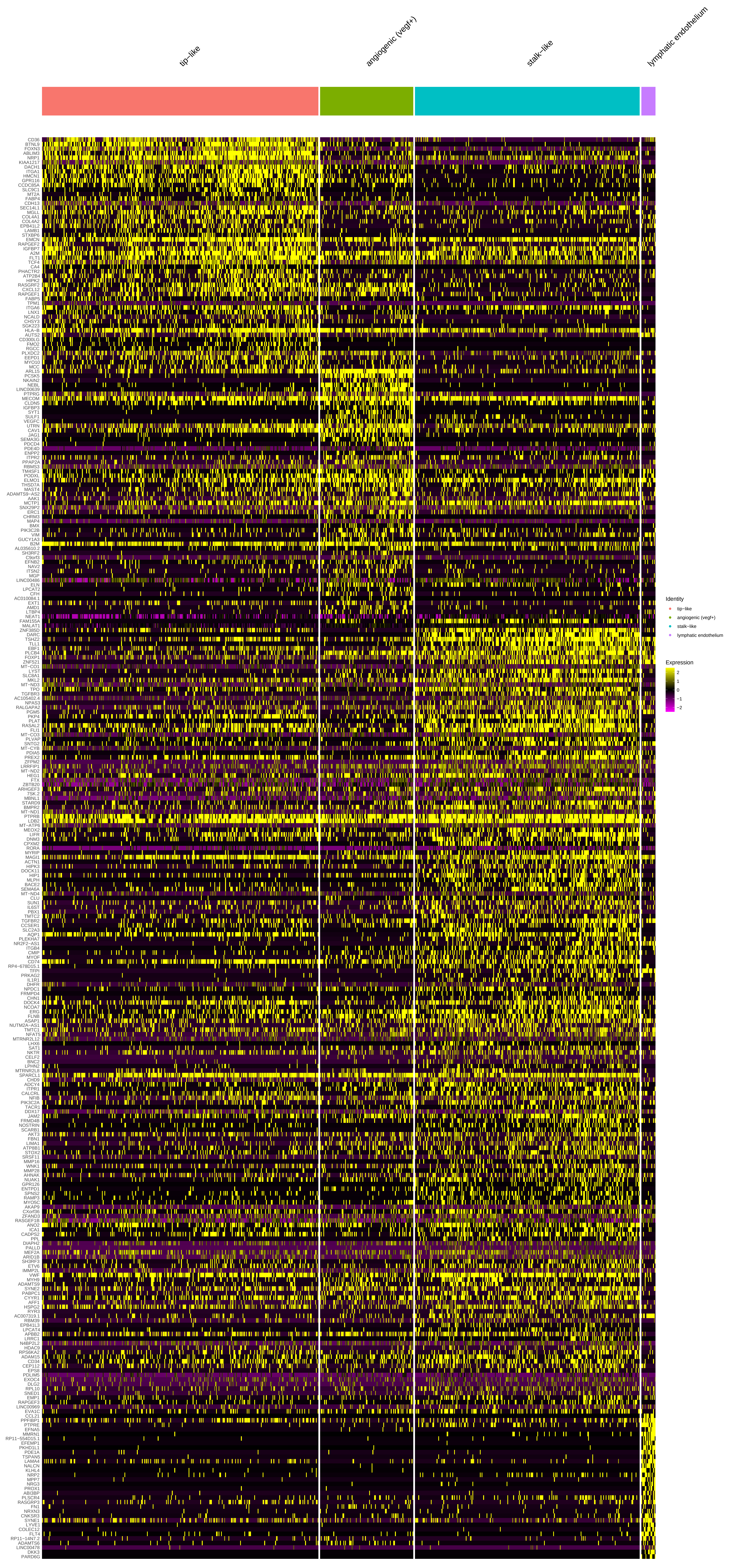

### d. Human Smooth Muscle heatmap

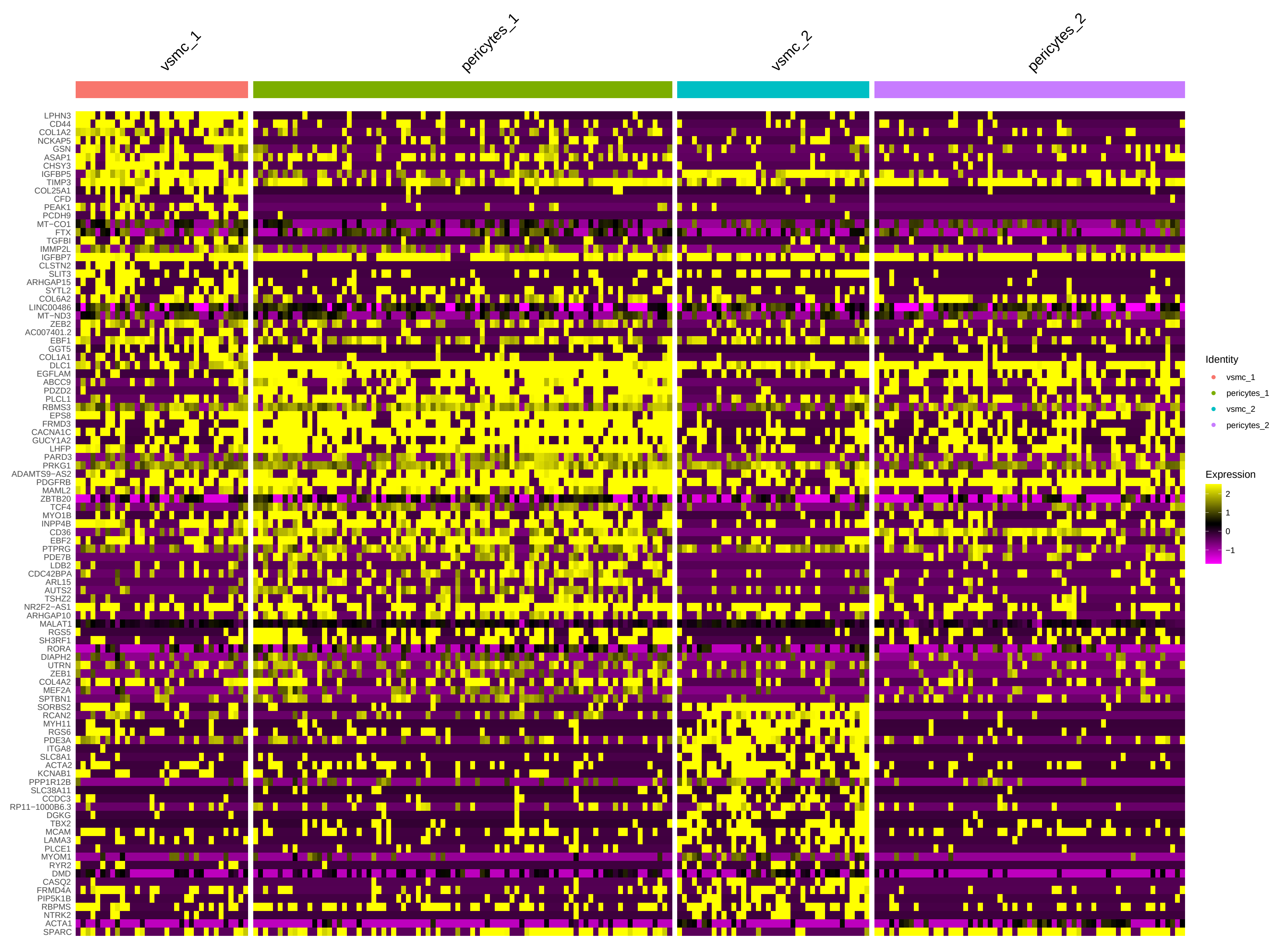


### e. Human Macrophage heatmap

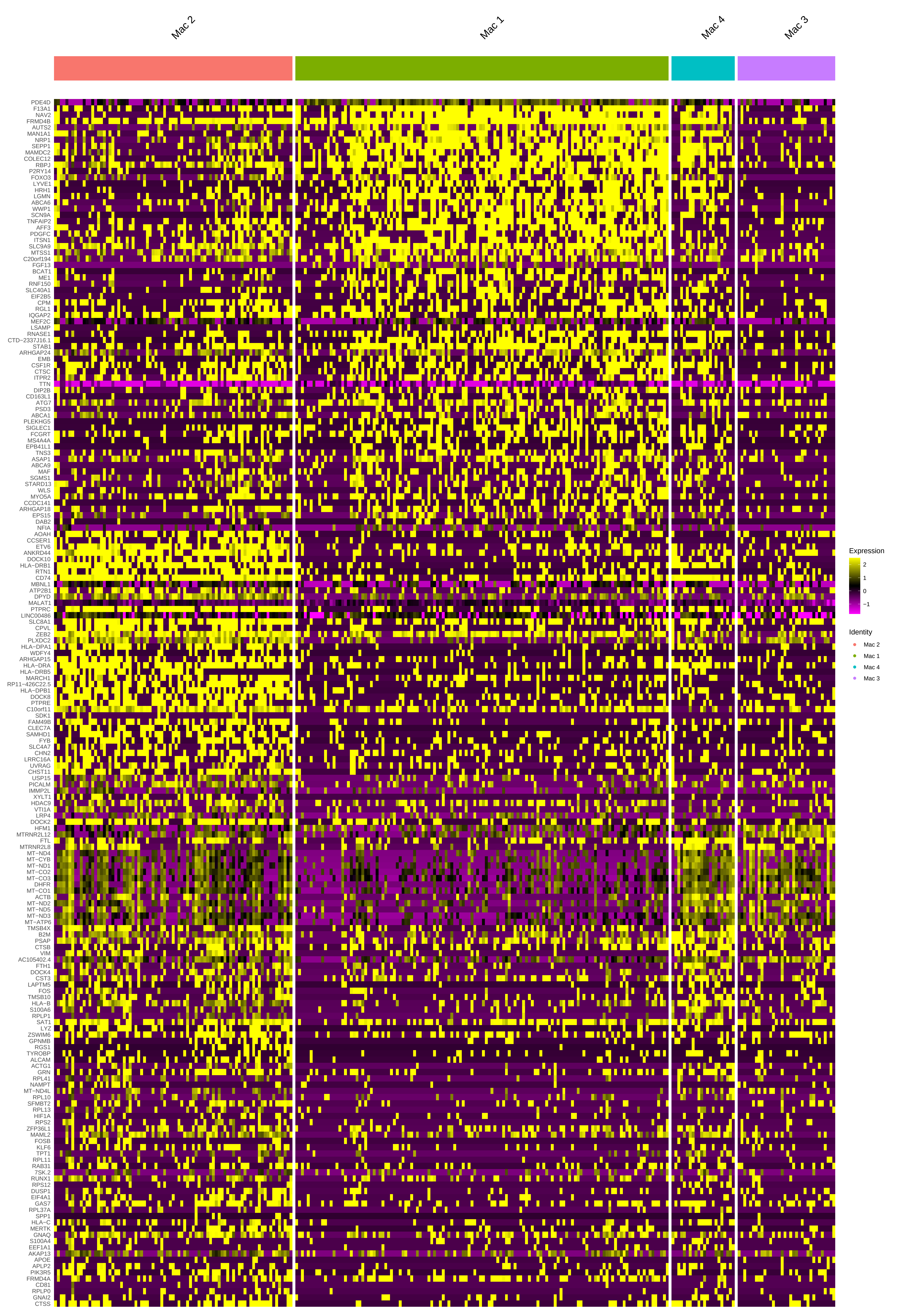
