## Supplemental Figures and References for "Single nuclei transcriptomics of muscle reveals intra-muscular cell dynamics linked to dystrophin loss and rescue"

**Supplementary Materials:**

Supplemental references: 65 - 248

Figure S1: Lineage tracing analysis of subpopulations identifies functional relatedness of endothelial and smooth muscle sub populations

Figure S2: heat maps mouse: A-z of all heatmaps not shown in main text

Figure S3: Dystrophin staining of whole murine quadricep from treated and untreated *mdx* mice

Figure S4: e23AON treatment in *mdx* shifts multiple distinct cell types towards more WT behavior (additional cell types)

Figure S5: heat maps human: A-z of all heat maps

Table S1: Key genes and references

Data file S1: Seurat findallmarkers files for mouse data

Data file S2: MDSC vs. M2MDSC differentially expressed genes

Data file S3: Differentially Expressed genes *mdx* vs *mdx* e23AON vs WT mouse data

Data file S4: MDSC vs. M1/M2 transitional differentially expressed genes

Data file S5: Fb POSTN1 vs. Fb CCL11 transitional differentially expressed genes

Data file S6: Seurat findallmarkers files for human data

Data file S7: Differentially expressed human genes; DMD vs healthy control data

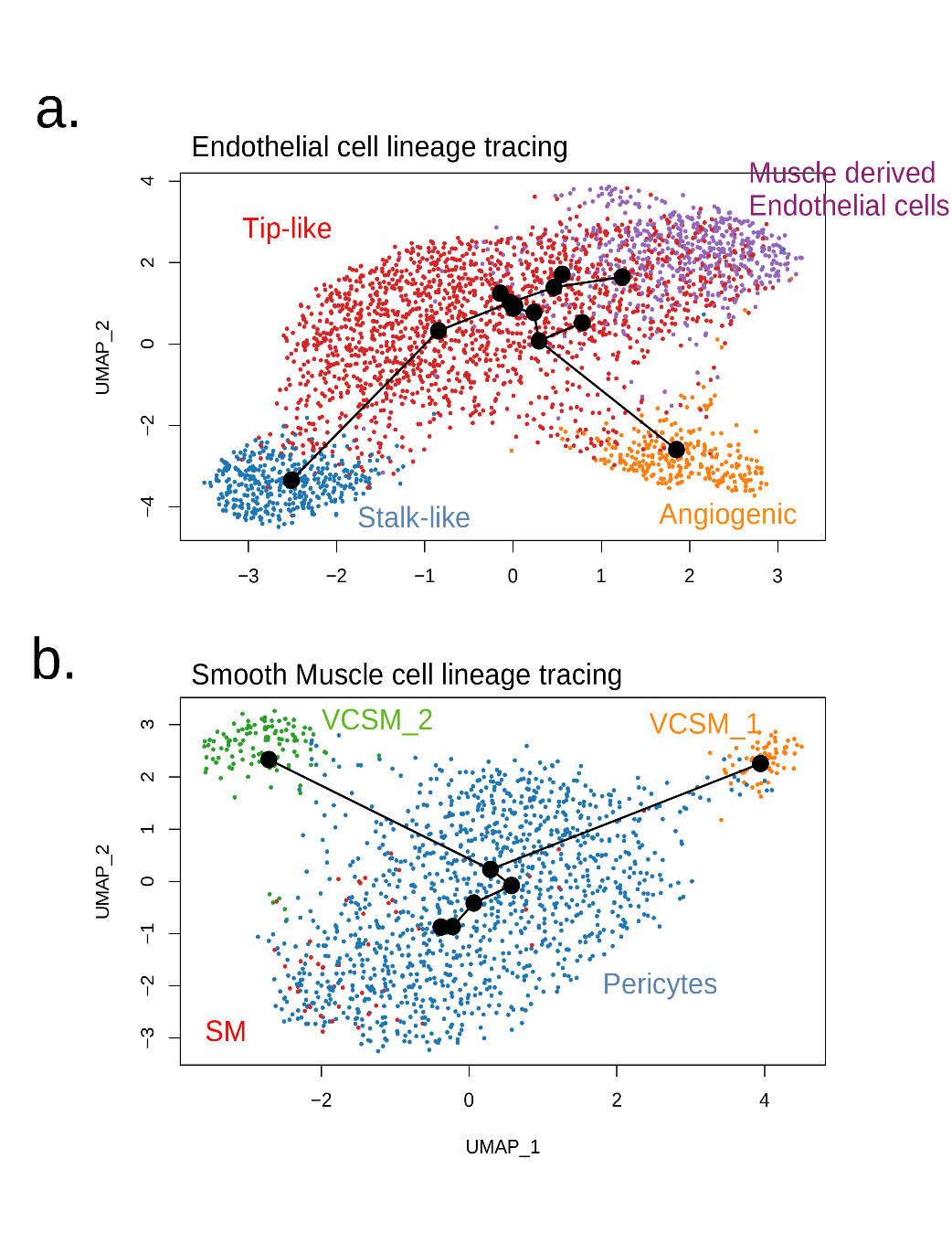

**Fig. S1**. Lineage tracing analysis of subpopulations identifies functional relatedness of endothelial and smooth muscle sub populations.

Lineage tracing analysis of (**a**) Endothelial subpopulations, (**b**) Smooth muscle subpopulations. Text in plots mark each subcluster population adjacent to each population and colored to match the sub-cluster identity of panel Fig. 1c.

**Too large for word include in document, see “Supp fig 2.pdf”**

**Fig. S2**: heat maps mouse: all heatmaps not shown in main text

Heatmap of significant marker genes among related mouse intra muscular cell types. All significant marker genes (Datafile S1) were used to generate these heatmaps, gene names listed on the beginning of each row. Each column represents an individual cell (grouped together by cell type identification). (**a**) Myofibers, (**b**) Satellite cells and Myoblasts, (**c**) Immune cells, (**d**) Fibroblasts/FAP, (**e**) Endothelial cells, (**f**) Smooth Muscle.

**Fig. S3.** Dystrophin staining of whole murine quadricep from treated and untreated *mdx* mice

### a. MOUSE 25- Untreated

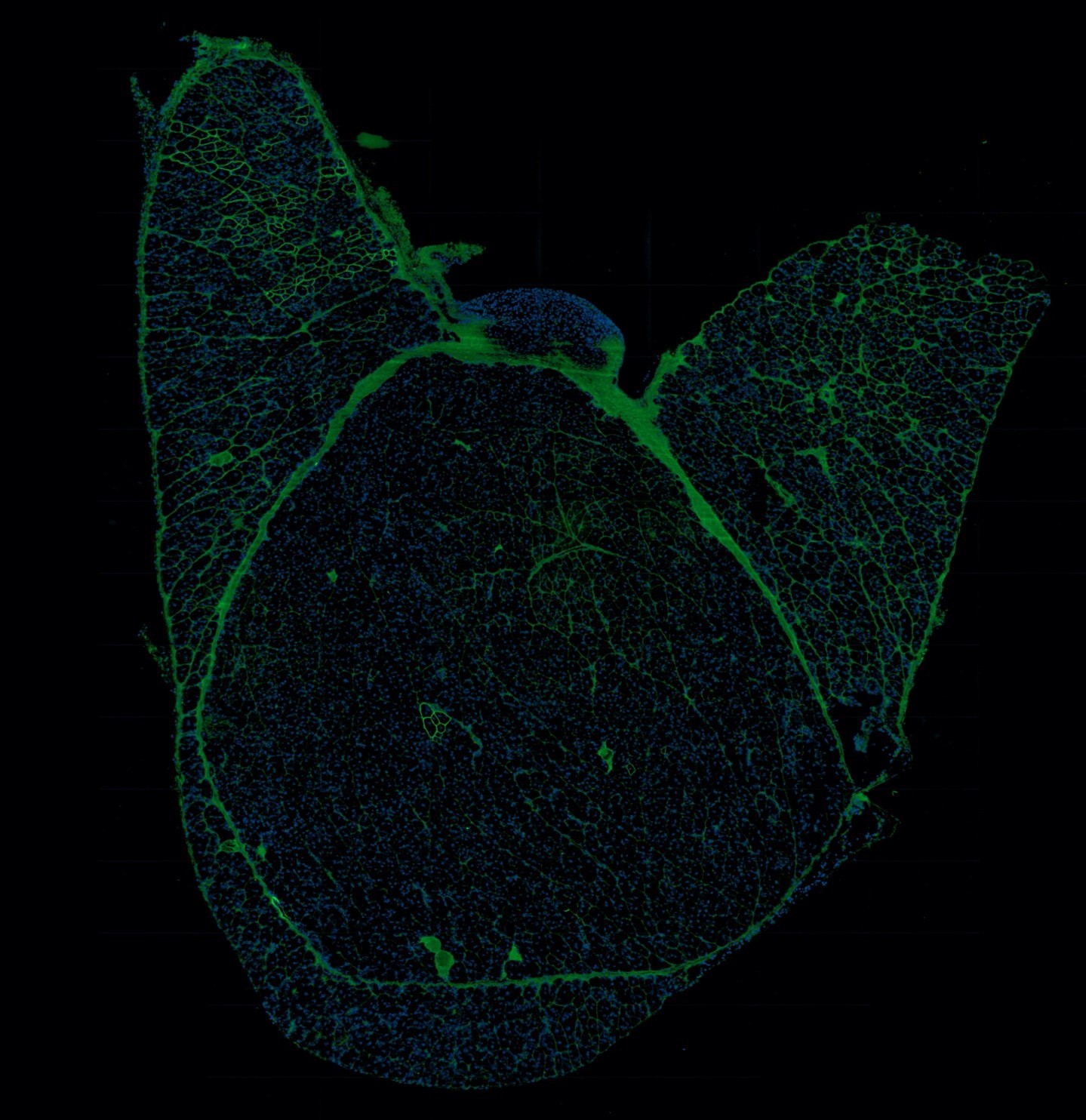

b. MOUSE 73- Untreated

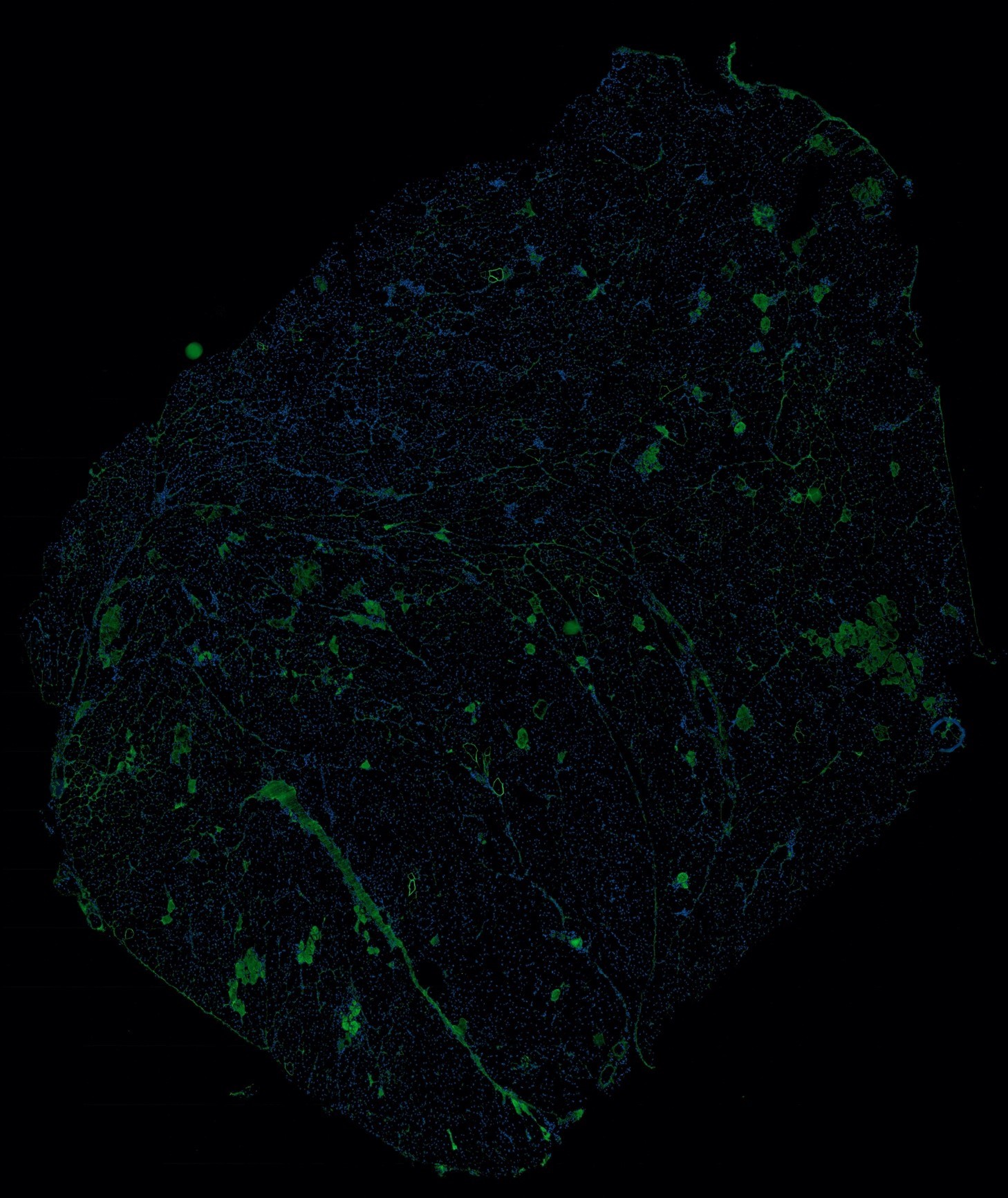

c. MOUSE 128- Untreated

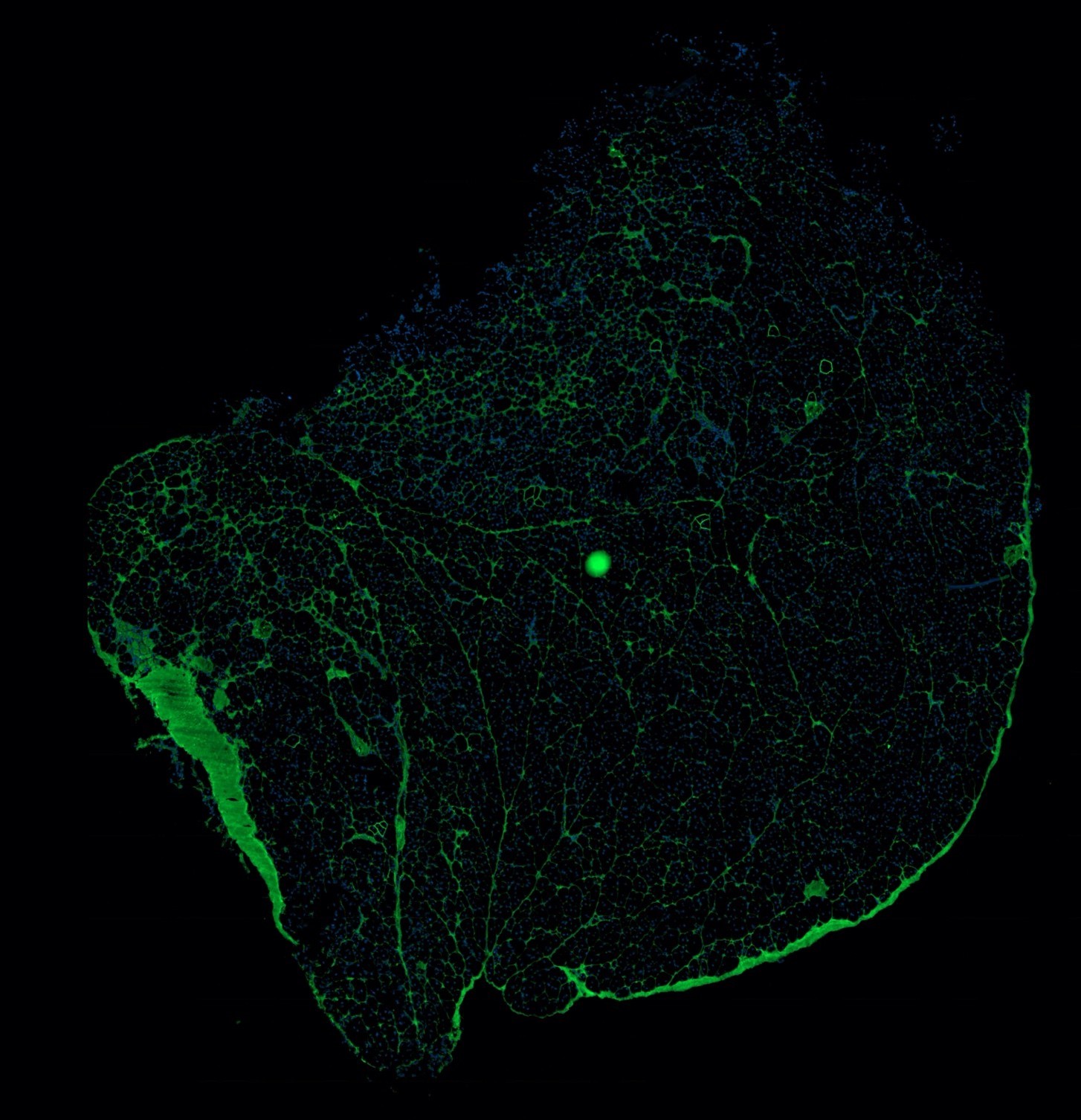

d. MOUSE - 155 Untreated

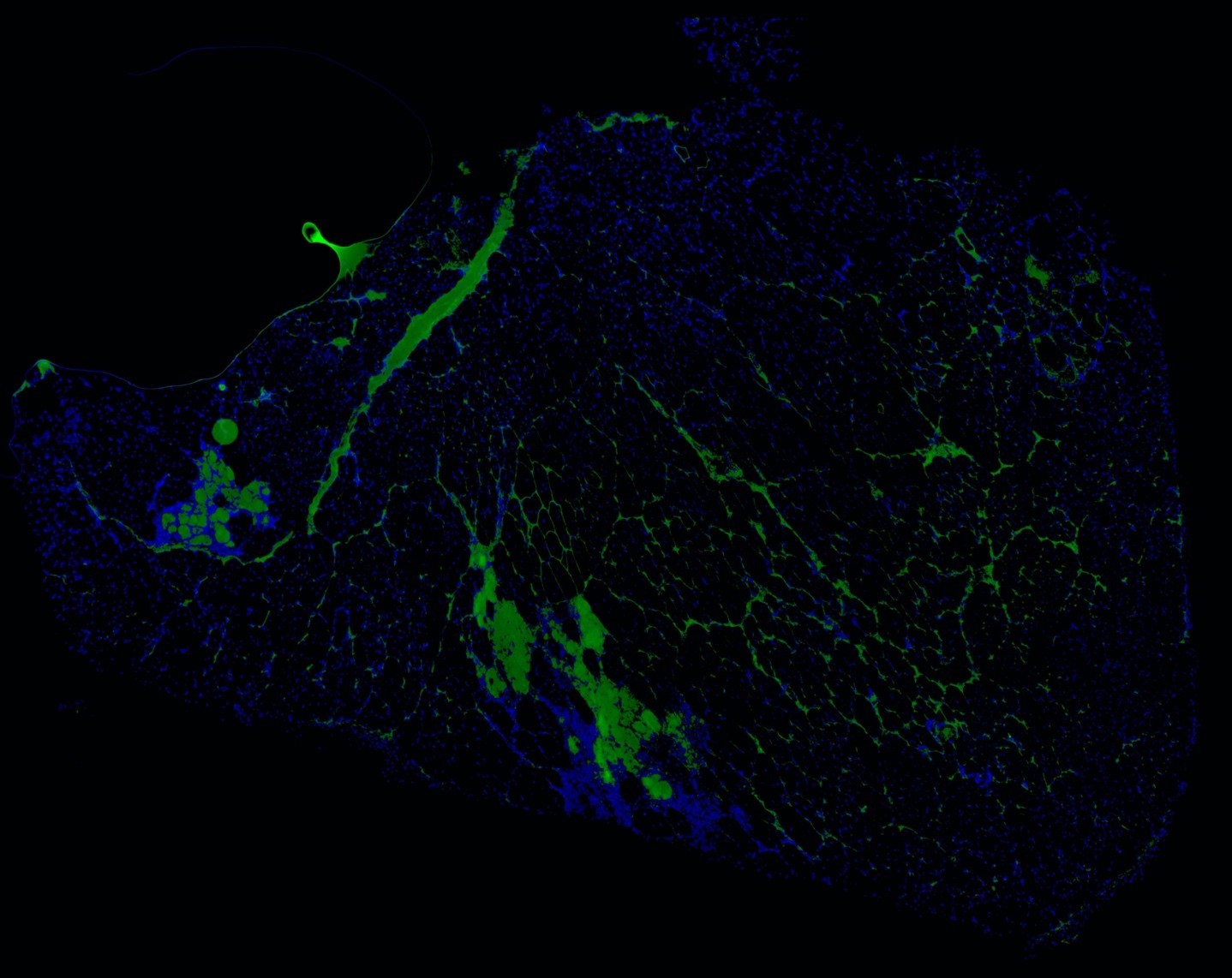

e. MOUSE 43- e23AON Treated

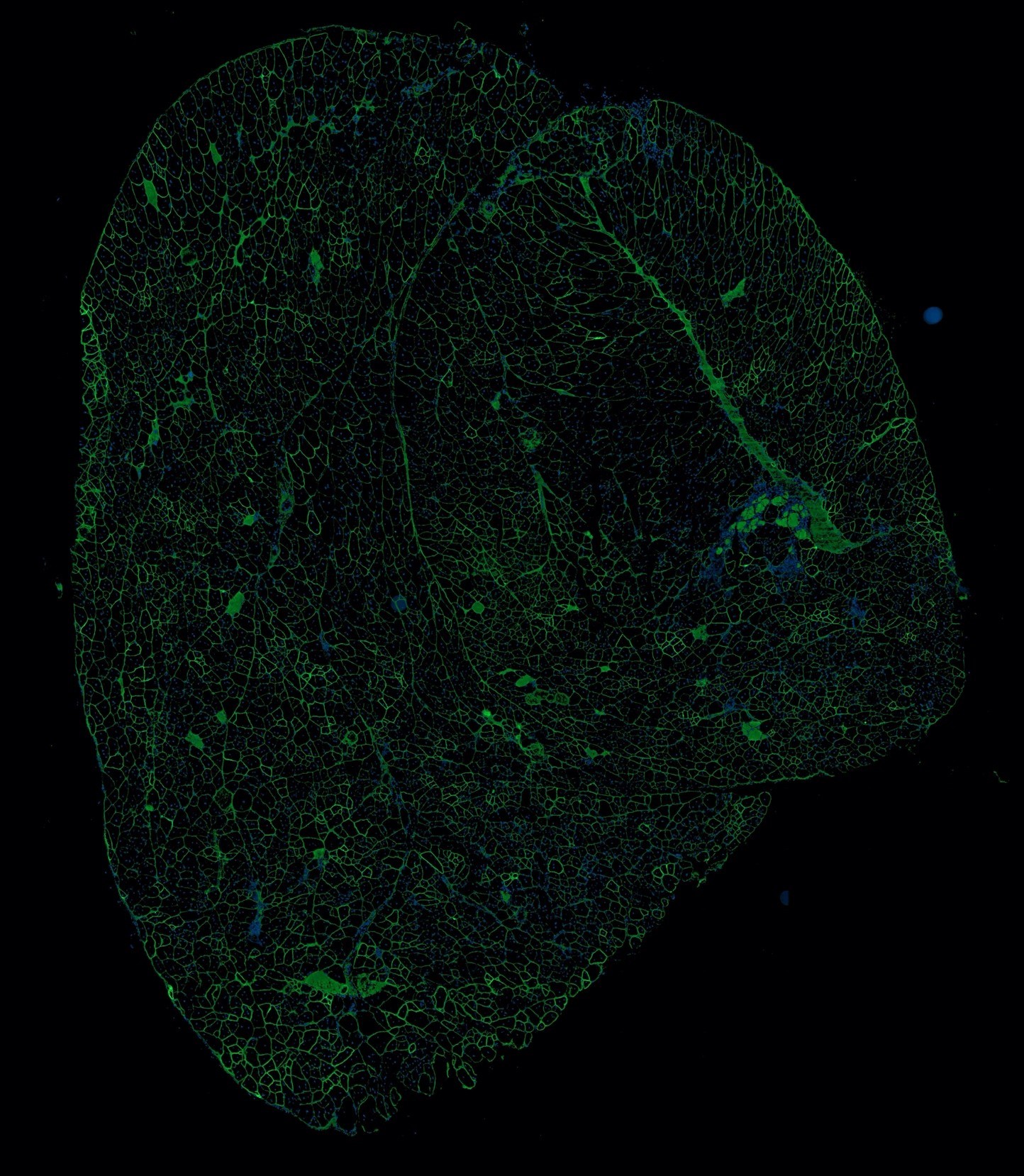

f. MOUSE 45- e23AON Treated

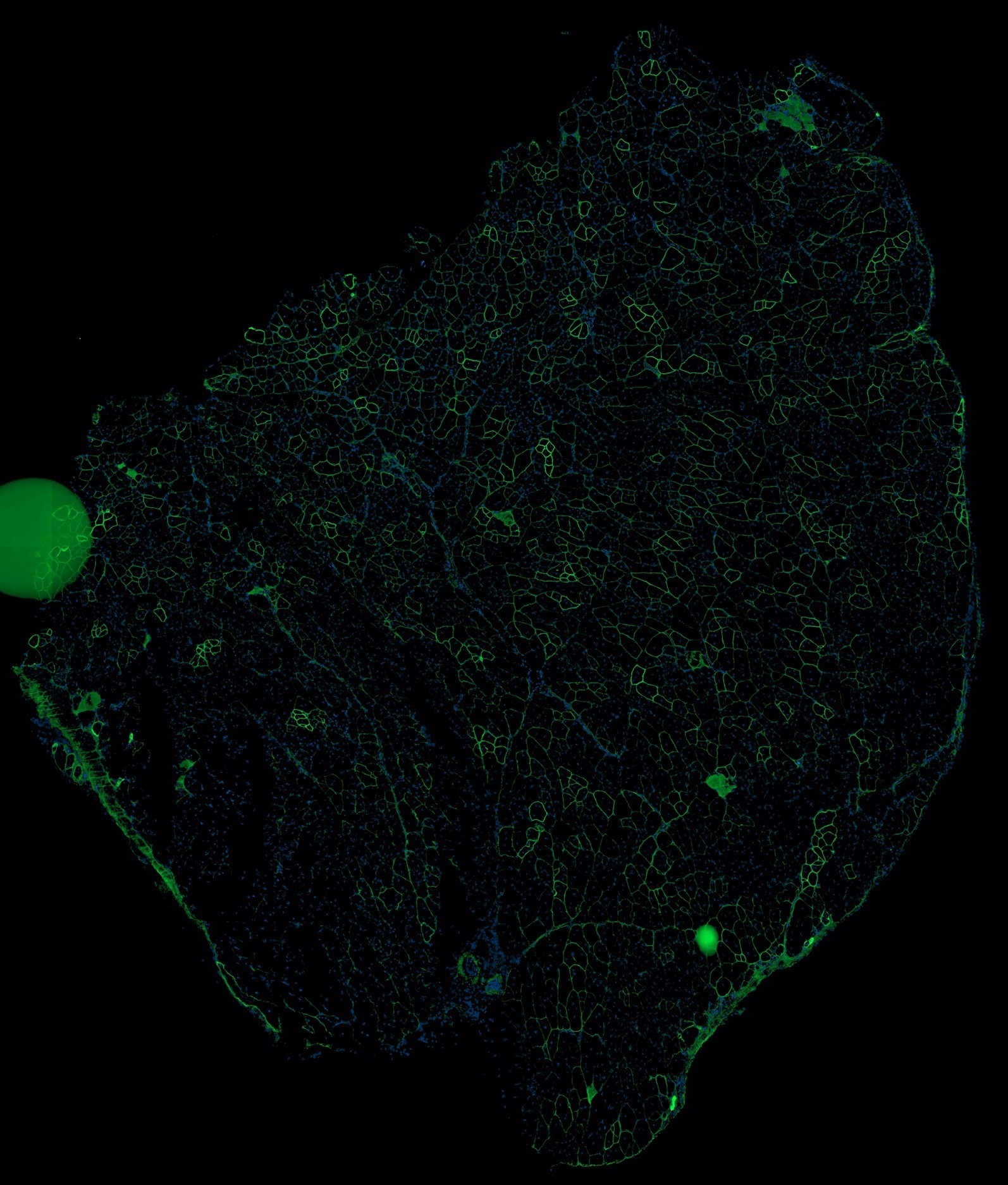

g. MOUSE 152- e23AON Treated

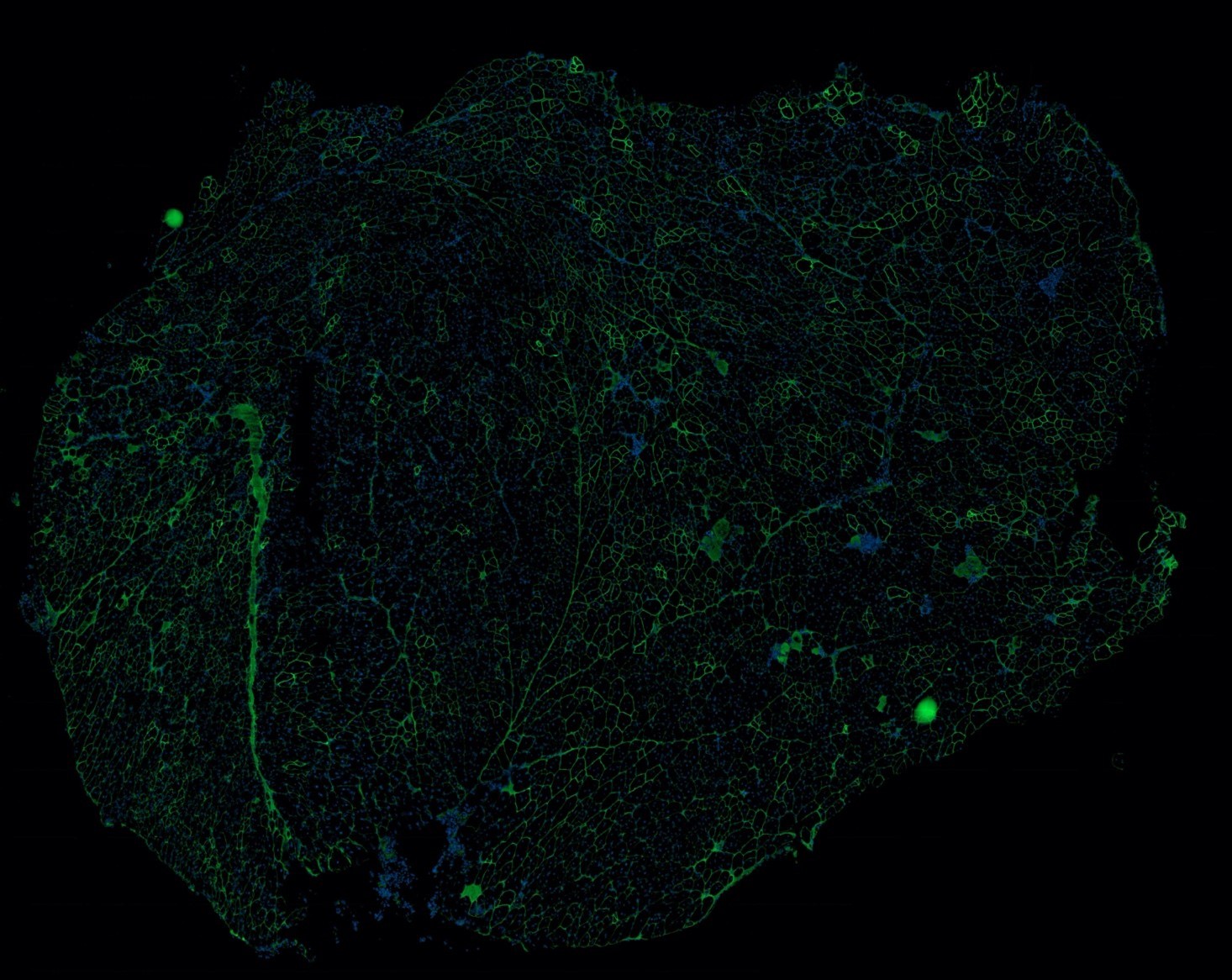

h. MOUSE 169- e23AON Treated

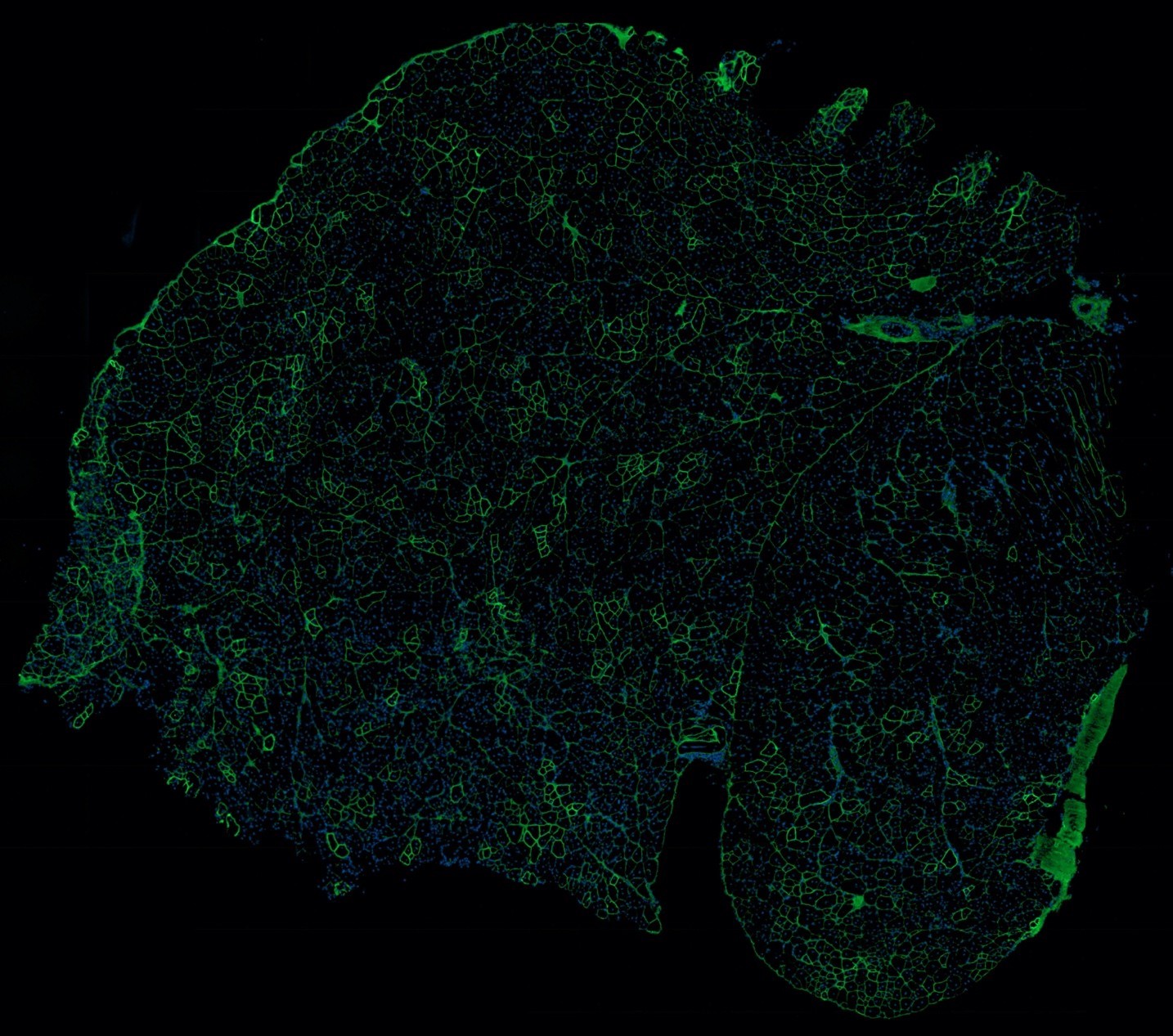

i. MOUSE C57- Wild type/control mouse

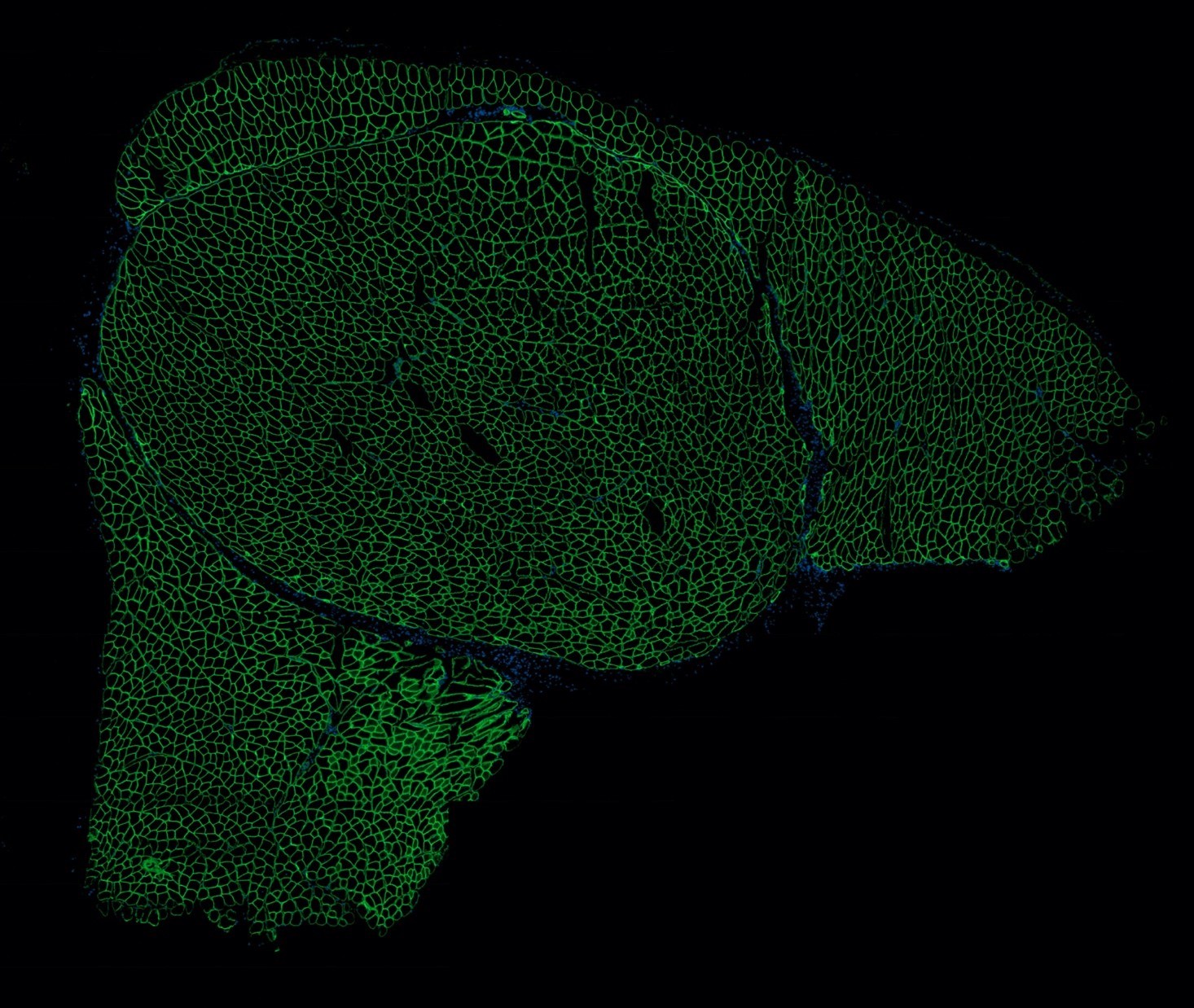

Fig. S3 Whole quadricep muscles from untreated *mdx* (**a** - **d**), *mdx* e23AON treated (**e** - **h**), or wildtype mice (**i**) were frozen, cryosectioned and subjected to immunofluorescent staining for dystrophin protein (green).

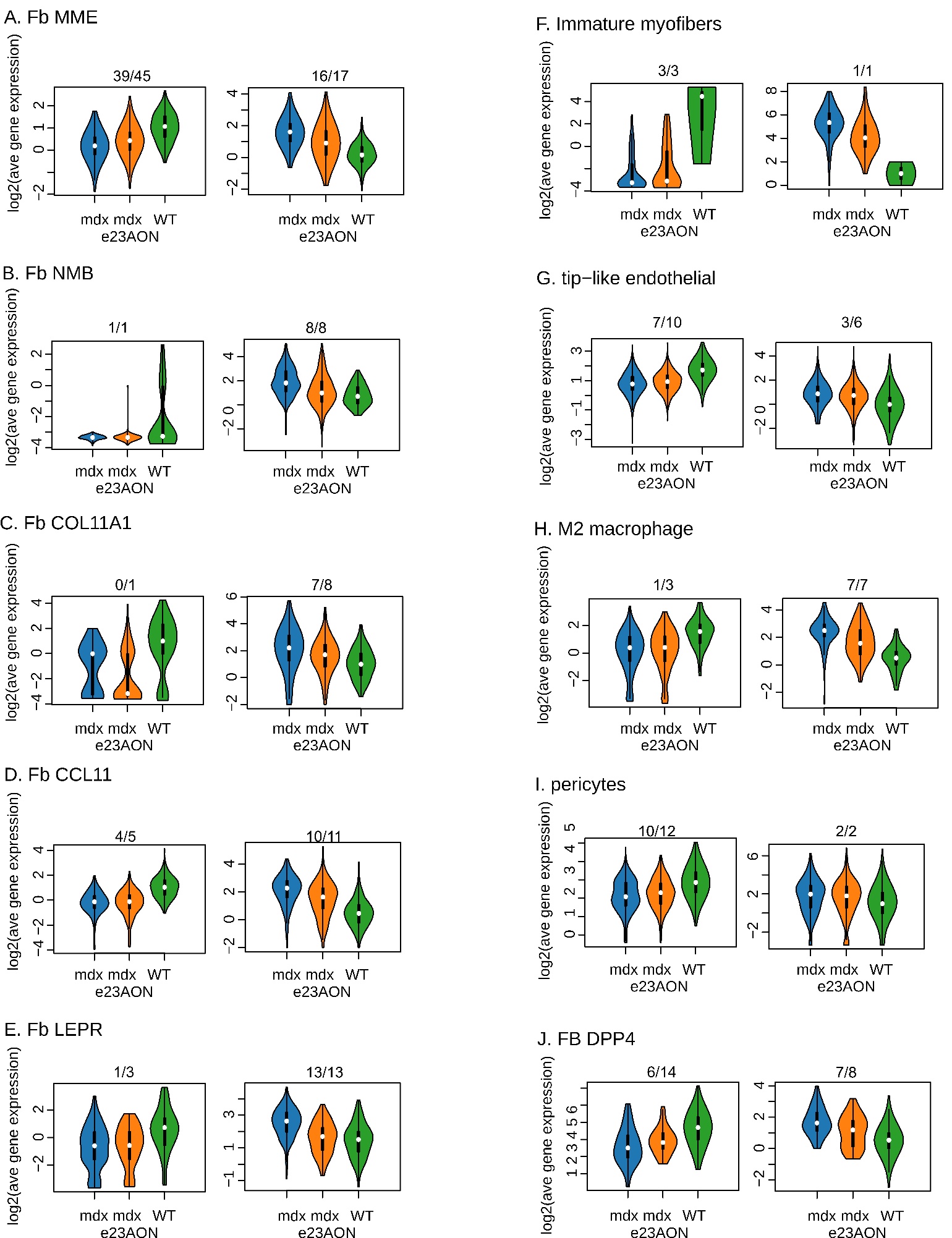

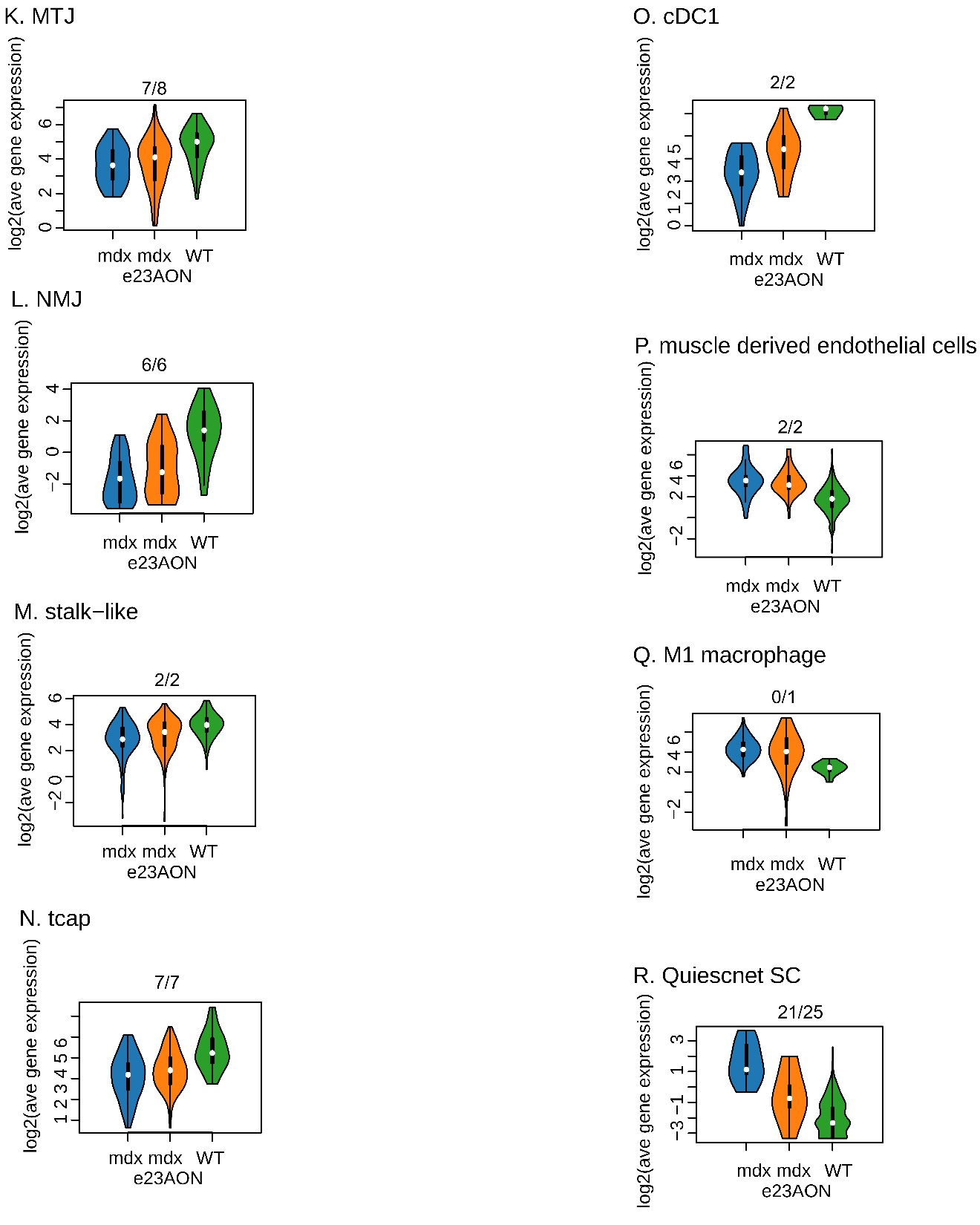

**Fig. S4** e23AON treatment in *mdx* shifts multiple distinct cell types towards more WT behavior (additional cell types)

Similar to Fig 5A. Average expression of genes upregulated (**left**) or downregulated (**right**) in WT relative to untreated *mdx*. Cell type names are listed above plot. Y axes = log_2_ average UMI for differentially expressed genes. (**l** - **p**) no significant genes were detected to be down regulated in WT. (**q** – **s**) no significant genes were detected to be up regulated in WT.

**Too large for word include in document, see “Supp fig 5.pdf”**

**Figure S5**: heat maps human: all heatmaps not shown in main text

Heatmap of significant marker genes among related human intermuscular cell types. All significant marker genes (Datafile S1) were used to generate these heatmaps, gene names listed on the beginning of each row. Each column represents an individual cell (grouped together by cell type identification). (**a**) Myofibers, (**b**) Satellite cells and Myoblasts, (**c**) Immune cells, (**d**) Fibroblasts/FAP, (**e**) Endothelial cells, (**f**) Smooth Muscle.

Table S1: Genes and References

**Table too large for word document, see “Table S1 – Genes and References.xlsx”**
